# Direct observation of the mechanical role of bacterial chaperones in protein folding

**DOI:** 10.1101/2020.10.20.346973

**Authors:** Deep Chaudhuri, Souradeep Banerjee, Soham Chakraborty, Shubhasis Haldar

## Abstract

Protein folding under force is an integral source of generating mechanical energy in various cellular processes, ranging from protein translation to degradation. Although chaperones are well known to interact with proteins under mechanical force, how they respond to force and control cellular energetics remains unknown.

To address this question, we introduce novel real-time magnetic-tweezers technology to mimic physiological force environment on client proteins, keeping the chaperones unperturbed. We studied two structurally distinct client proteins with seven different chaperones, independently and in combination, and proposed novel mechanical activity of chaperones. We found chaperones behave differently, while these client proteins are under force than its previously known functions. For instance, tunnel associated chaperones (DsbA and trigger factor), otherwise working as holdase without force, assist folding under force. This process generates an additional mechanical energy up to ∼147 zJ to facilitate translation or translocation. However, well-known cytoplasmic foldase chaperones (PDI, thioredoxin, or DnaKJE), does not possess the mechanical folding ability under force. Notably, the transferring chaperones (DnaK, DnaJ, SecB), act as unfoldase and slow down folding process, both in the presence and absence of force, to prevent misfolding of the client proteins. This provides an emerging insight of mechanical roles of chaperones: they can generate or consume energy by shifting energy landscape of the client proteins towards folded or unfolded state; suggesting an evolutionary mechanism to minimize the energy consumption in various biological processes.

## Introduction

Protein folding under force is an important source of generating mechanical energy that harnesses different biological functions ranging from protein translation to degradation^1–9^. For example, in translation, protein synthesis occurs in a confined ribosomal tunnel under 7-11 pN force while ClpX machinery generates 4-15 pN force to unfold protein followed by ClpP dependent degradation^2, 6^. Notably, molecular chaperones, well involved in various folding process^10–14^, must interact with proteins under force. Recent force spectroscopy studies have nicely illustrated the molecular mechanisms of various chaperones, but most of them focused on the role of chaperones on the folding-misfolding aspects of the client proteins^15–17^. However, few studies reported an emerging evidence that molecular chaperones could modulate the force transmission through their client polypeptide during different folding-associated processes including, during co-translational folding at ribosomal tunnel edge^6^. For instance, trigger factor (TF), a ribosomal tunnel-associated chaperone, increases the transmission force of the nascent polypeptide chain, by assisting their folding under higher force^18,19^. Similarly, a prototypical chaperonin GroEL mediates the forced unfolding of misfolded proteins on their apical domain^20,21^. Therefore, the generation of mechanical energy during these diverse chaperone-protein interactions under force could be a linchpin factor in understanding the biological system. Furthermore, the mechanical activity by all of these molecular chaperones indicates a broader mechanical role of chaperones than previously assumed, but how these chaperones modulate the mechanics of protein folding at the molecular level remains largely unknown.

To address these questions, we investigated the mechanical role of different chaperones and chaperone-like proteins under force, individually and in combinations: DnaK, DnaJ, SecB, DsbA, Trigger factor (TF), protein disulfide isomerase (PDI), and thioredoxin (Trx). The mechanical effects of these chaperones were monitored by our novel real-time magnetic-tweezers that implements both force constant force and force ramp methodology^22^. Using the constant force methodology, a constant force can be provided on a protein, offering a unique way to probe both thermodynamics and kinetic properties of the substrate protein under equilibrium conditions. Whereas, the force ramp methodology, by increasing or decreasing the force linearly at a loading rate, detects unfolding and refolding events at different forces.

Furthermore, the advantage of broad force ranges from 0 to 120 pN, with a sub-pN resolution, allows us to monitor the effect of chaperone on both the folded and unfolded states of the substrate independently. Moreover, the associated fluid chamber offers to introduce different chaperones, one by one or in combination, and tracking their dynamic changes on a single client protein molecule in real-time. Lastly, using this technology, the force can be precisely applied on the substrate protein while keeping the chaperone unperturbed.

We have systematically explored the mechanical activity of different chaperones on two structurally distinct substrates i.e., protein L which predominantly contains β sheets and β-hairpins in their structure and talin containing four α helices packed against a buried hydrophobic threonine belt. Although these chaperones are structurally different, we found they function similarly with chaperones. Among these chaperones, proteinL has been used as chaperone substrate previously^17^, while talin being a ‘large’ cytoplasmic protein could potentially interact with any cytoplasmic chaperones.

Our results showed that chaperones behave differently while the substrate is under force than their known role. Prototypical chaperone TF, having a well-known holdase function in the absence of force^19,23^, behaves differently, while the substrate protein is under force, by accelerating refolding kinetics and generates a mechanical work of 147 zJ. A distinct property is also observed for DnaKJE (DnaK+DnaJ+GrpE+ATP) complex under force. In the absence of force, this complex work as a typical foldase^24, 25^; but in the presence of force, it does not show its intrinsic foldase activity. Furthermore, oxidoreductase enzyme DsbA, which has been observed to possess ‘weaker chaperone’ activity than cytosolic oxidoreductase PDI in the absence of force^26^, exhibits stronger foldase activity than PDI and Trx under mechanical force. However, DnaK, DnaJ, SecB chaperones, both in the absence and presence of force, function as holdase and reduce the mechanical work by hindering the folding process. Thus, based on our observations with two structurally distinct substrates (protein L and talin), we reclassified the chaperones into three classes on the basis of their mechanical characteristics: mechanically unfoldase (DnaJ, DnaK, and SecB); mechanically neutral (DnaKJE, PDI, and Trx), and lastly, mechanically foldase (TF and DsbA). Chaperones associated with tunnels, TF and DsbA possess a special mechanical activity under force by which they can generate extra mechanical power to pull protein from ribosome to cytoplasm and cytoplasm to periplasm. By contrast, the same protein, in the absence of force, behaves as a holdase^19, 23^. Overall, these data reconcile that chaperone, by modifying free energy landscape of substrate proteins, generates the mechanical energy that provides a distinct view of mechanical folding pathway of proteins under equilibrium.

## Results

### Single-molecule measurements of protein folding by real-time magnetic tweezers

Real-time magnetic tweezers allow us to probe the folding-unfolding dynamics of a single protein in presence of a chaperone where the force can only be applied on the proteins while chaperones remain unaffected. Additionally, the force can be precisely controlled in sub-pN resolution that allows us to observe the dynamic folding-unfolding transition in the substrate polyprotein, providing information at equilibrium force. We performed the experiment on an engineered construct containing eight repeats of protein L B1 domain and monomer of talin R3-IVVI domain, inserted between N terminus HaloTag and C terminus AviTag. The proteins are tethered on a glass surface by covalent HaloTag chemistry and attached to the paramagnetic beads by biotin streptavidin-biotin chemistry. Applied force on paramagnetic beads is regulated by a pair of permanent magnets, attached to the voice coil actuator. The details of force calibration can be found in the supplemental information (Supplementary Figure 10-12).^22, 27, 28^ In short, the force magnitude was calibrated by changing the magnet-bead distance such as moving down the magnet position from 4.0 mm to 1.4 mm results in an increase in force pulse from 4.3 pN to 45 pN (Fig. 1A). At unfolding pulse of 45 pN, the polyprotein undergoes an elastic extension, followed by the unfolding of tandem domains, which are observed as eight unambiguous stepwise extension of ∼ 15 nm (described as the difference in length between folded and unfolded domain).

**Figure 1:**
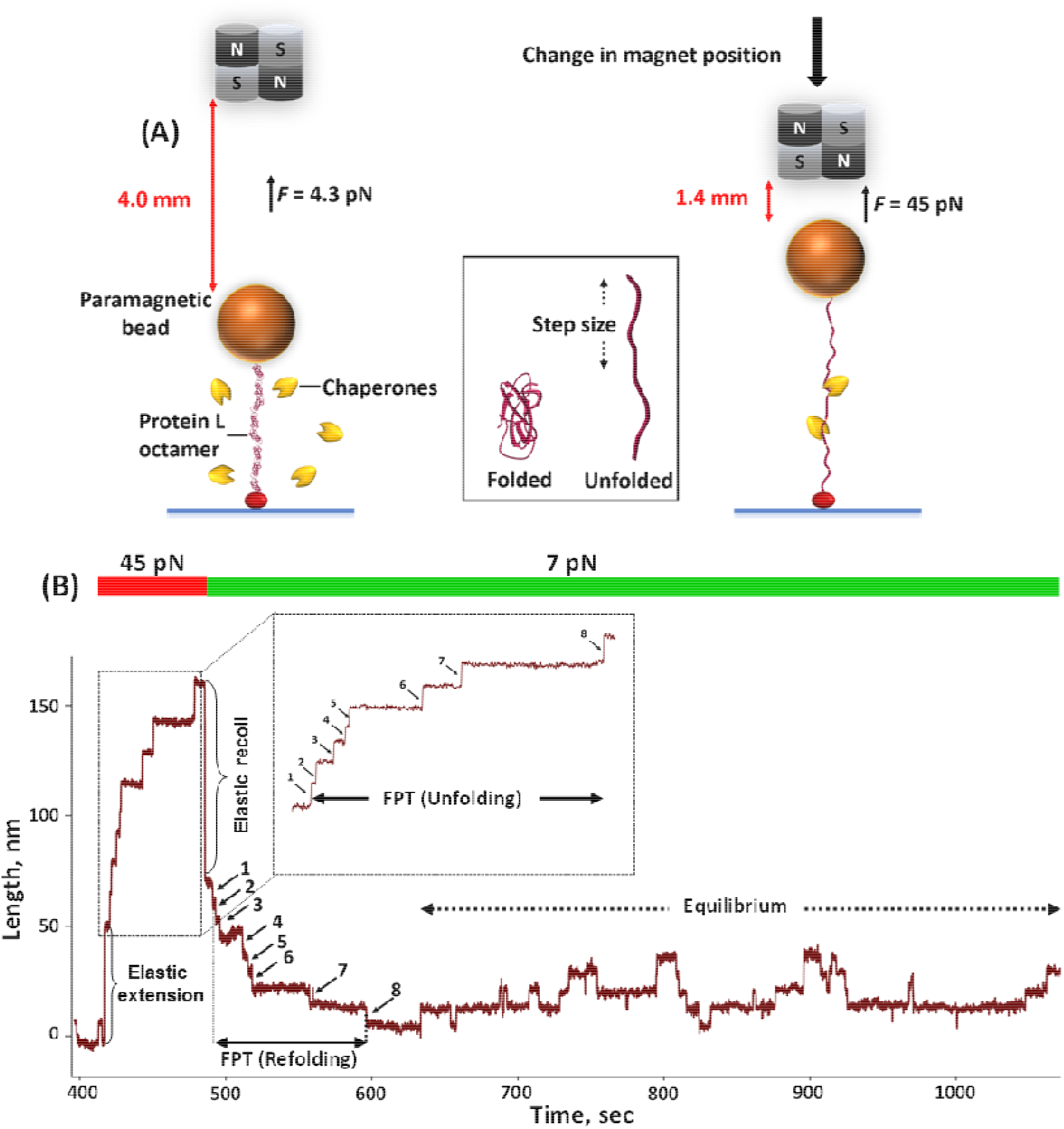
Experimental set-up for real-time magnetic tweezers to study protein-chaperone interaction: **(A) Schematics of magnetic tweezers experiment:** The engineered construct of eight identical protein L domain, inserted between N terminus HaloTag for attaching to ligand coated glass surface and C terminus AviTag for binding to streptavidin coated paramagnetic beads. Applied force is controlled by changing the distance between permanent magnet and paramagnetic beads. The distance between folded and unfolded state is denoted as step-size. **(B) Standard folding trajectory of protein L octamer:** First, the polyprotein is unfolded at high force pulse of 45 pN, generating eight successive steps as single molecule fingerprint that followed by a refolding pulse at 7 pN. During refolding, the polyprotein first undergoes a relaxation phase followed by an equilibrium phase where the domains show a continuous hopping between folded and unfolded state. The minimum duration of complete unfolding of all the domains in the polyprotein from fully folded state is called first passage time (FPT) of unfolding and the time taken to achieve complete refolding of eight domains from the fully unfolded state is called FPT of refolding.

Fig. 1B demonstrates the representative folding trajectory obtained by our single molecule experiment that consists of two phases: at first, the polyprotein is unfolded at 45 pN to generate eight discrete stepwise extensions as a single molecule fingerprint and subsequently quenched to a refolding pulse of 7 pN. During refolding, the polyprotein first experiences elastic recoiling and a relaxation phase (non-equilibrium phase), observed as a downward refolding staircase and is characterized for kinetics by mean-first passage time (MFPT) for fully folded state and similarly during unfolding, MFPT for fully unfolded state can be determined for the unfolding kinetics^19, 29^. This phase is followed by an equilibrium phase where the domains exhibit folding-unfolding dynamics. The folding-unfolding transitions are monitored as downward steps of refolding and upward steps of unfolding, suggesting a balanced condition for the substrate protein. The unfolding trajectory is magnified in the inset of Fig. 1B where the FPT of unfolding is denoted as-minimum time taken for complete unfolding of eight domains from fully folded state and similarly, for FPT of refolding, time taken to complete refolding of all domains from the fully unfolded state. Mean-FPT (MFPT) is calculated by averaging the FPT values over numerous trajectories, providing a model-free metric that describes unfolding and refolding kinetics of proteins under force^19, 29^.

We determined the folding probability (FP) by dwell-time analysis, exclusively from the equilibrium phases over many folding trajectories, as described previously (Fig. 1B)^7, 19, 30, 31^. The relative population of folded and unfolded states at equilibrium phase is strongly dependent on applied force. For instance, at 4 pN, the FP∼1 while at 12 pN, it decreases to zero (FP∼0). We measured the folding probability in the narrow force range within 4-11 pN where both folding and unfolding transitions can be observed over an experimental time-scale and fitted to the sigmoidal equation. Along with the equilibrium phase, each folded state is named as *I* (number of folded states) and their relative population at a certain force is determined as *t*= *T_I_*/*T_t_*, where *T_I_* is the cumulative duration of *I* state and *T_I_*. = ∑*_I_ T_I_*, total observable duration of the equilibrium phase (Supplementary table 1). FP is calculated as a normalized averaged state (Supplementary Table 1)^7, 19, 30, 31^,

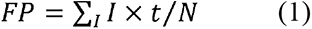

Where N is defined as the numbers of total domains. Thus, in a simple sense, the folding probability can be calculated as a ratio of refolded domains to the total number of domains in the polyprotein construct. To confirm that the domains are properly refolded, we extended the substrate protein at high force (probe pulse) at the end of the experiment. The number of unfolded steps is matching exactly with the number of domains folded, when the probe pulse is applied (Supplementary Figure 1). Detailed calculations of folding probability have been explained in the supplementary information, whereas the free energy of protein folding under equilibrium can be described as follows^32^.

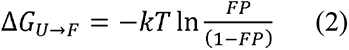

### Protein L does not form misfolded state under force

The chaperone mechanisms are mostly studied with the substrate proteins, which are prone to form misfolded state and aggregation^16, 33–36^. To eliminate diverse conformational dynamics, arising from protein misfolding, we used well-known protein L B1 domain that does not form misfolded state and studied as chaperone substrate previously^19, 22, 37^. B1 domain is an IgG-binding domain, containing 62 residues with no proline, forms a globular fold conformation. The domain structure includes a four-stranded beta sheet, packed against a central α-helix with two β-hairpin motifs. The first β-hairpin contains a turn while the second hairpin has conformational strain due to consecutive positive φ angles in backbone^38^. To show this protein L domain does not form any misfolded state, under our experimental condition, we used a continuous force ramp protocol on protein L. Force ramp is an excellent methodology to monitor misfolded states, which are mechanically weak and thus, unfold at lower force^39^. We run successive ten unfolding-refolding cycles of a linear force-increase scan from 4 to 80 pN with a loading rate of 2.5 pN/s, followed by a force-decrease scan with the same loading rate. We applied the high force of 80 pN so that, all the eight unfolding events, occurred below this force, could be recorded. If protein L does not refold properly in the refolding pulse and form misfolded state, in the next unfolding pulse it will unfold at lower force and could be detected in the next unfolding pulse. In the force-decrease scan, the refolding events seems to be smooth due to the refolding of all the domains within the same force range and thus, no distinct refolding steps are observed. We monitored the average unfolding force in every pulse and plotted the average forces against pulse/cycle number (Supplementary Figure 2B). Our result shows the mechanical strength of protein does not change with the unfolding pulses, indicating the absence of any misfolded states (Supplementary Figure 2).

### Mechanical unfoldases induce mechanical unfolding and suppress refolding

DnaJ is a prototypical chaperone that holds the polypeptide from trigger factor and then, either transfers it to SecA translocon via SecB or folds with the help of DnaK/DnaJ/GrpE^15, 19, 40–42^. Chaperones, both in the absence and presence of force, are well-known to interact with the near-native or partially-folded, collapsed, and folded structures^15, 34, 43–48^; however, to further explore its effects on protein folding under force, we have measured unfolding and refolding kinetics, and folding probability of protein L. Fig. 2A describes the variation in MFPT of refolding as a function of force in both absence (black) and presence of 1 µM DnaJ (red). The MFPT of refolding steadily increases with force magnitude, however, at a particular force with DnaJ, the MFPT of refolding is higher than without DnaJ. For example, at 5 pN in absence of DnaJ, the polyprotein requires 6.3±2 s for complete refolding but with DnaJ, it increases to 25.4±0.2 s (Fig. 2A). This lower refolding kinetics data suggest that DnaJ interacts with the extended substrate under force. Similarly, unfolding MFPT has been observed to decrease steadily in the presence of DnaJ (Fig. 2B), signifying the destabilization of the collapsed state in the presence of DnaJ chaperone. To confirm whether the chaperones interact with the collapsed state of the protein substrate, we performed the force ramp experiment and observed that the mechanical stability of the protein is decreased by the unfoldase chaperones and increased by the foldase chaperones (Supplementary Figure 13).

**Figure 2:**
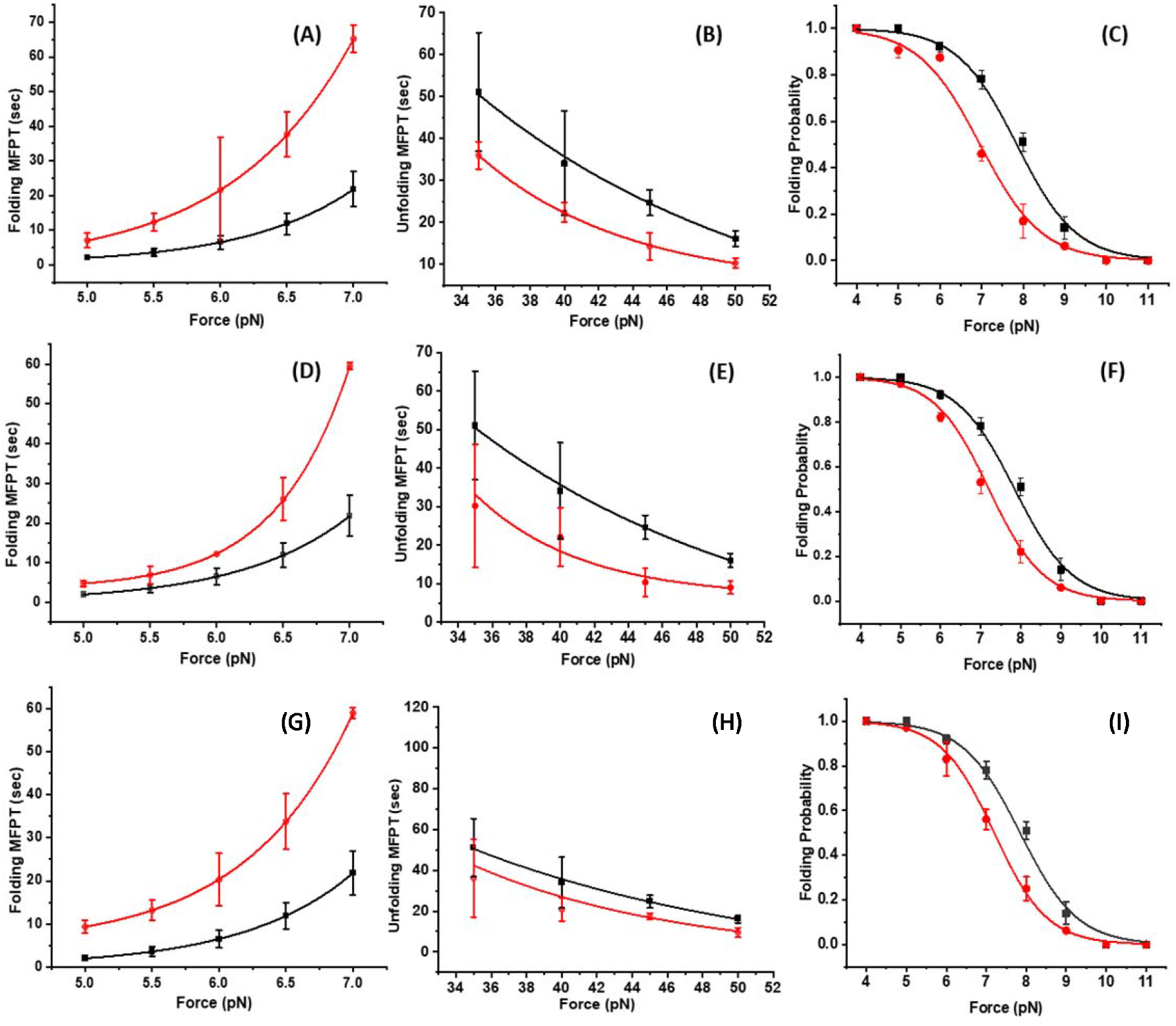
Unfoldase chaperones negatively modulate the folding events in protein L: Unfoldase chaperones hinders the protein folding by changing the MFPT values and it is plotted against the force in the absence (black) and presence of different chaperones (red). **(A)** DnaJ increases the MFPT of refolding (red) than the control (black), **(B)** while decreasing the MFPT of unfolding than in its absence, indicating faster unfolding kinetics with DnaJ. For each analysis, more than eight individual trajectories are measured and averaged. Error bars are standard error of the mean (s.e.m.). **(C)** Folding probability is plotted against the varying forces and DnaJ reduces the folding probability within 5-9 pN force range by shifting the half-point force from 8 to 6.9 pN, suggesting a pronounced reduction in folding probability in the presence of DnaJ. Data points are calculated using >2500s and over five molecules per force. Error bars are s.e.m. **(D)** DnaK, as a strong unfoldase, has also been observed to increase the MFPT of refolding (red) than the control (black), while **(E)** decreasing the MFPT unfolding (red) than the control (black). For unfolding and refolding MFPT analysis, ten and six individual trajectories are calculated, respectively. Error bars are s.e.m. **(F)** The presence of DnaK substantially decreases the folding probability at a particular force and thus, the half-point force is shifted from 8 to 7.2 pN. Data points are calculated using >2500s and over five molecules per force. Error bars are s.e.m. **(G, H)** Similarly, SecB increases the refolding MFPT and decreases unfolding MFPT than the control. More than eight trajectories are calculated for the MFPT analysis. Error bars are s.e.m. **(I) Folding properties:** SecB reduces the folding probability as a particular force and the half-point force is also reduced from 8 pN to 7.2 pN suggesting a 50% reduction in folding probability. Data points are calculated using >2500s and over five molecules per force. Error bars are s.e.m.

We also monitored the folding probability (FP) of protein L (control, black) and compared with that of in the presence of DnaJ (red). Folding probability decreases with increasing force, indicating an inverse relation of protein folding with force. We plotted FP as a function of force within 4-11 pN range and observed that at 4 pN, FP=1, irrespective to the presence of DnaJ and at 10 pN, it becomes zero. Interestingly, mechanical effect of DnaJ is prominent within 5-9 pN range, observed as the decrease in half-point force (defined as a force where FP is 0.5) from 8 to ∼6.9 pN (Fig. 2C); suggesting an optimum DnaJ binding to protein L which results in its lower refolding ability.

As a substrate presenter, DnaJ recruits the polypeptide to DnaK which acts as a holdase, reducing the refolding rate of its client protein^23, 33–36, 49^. Thus, to understand the effect of DnaK in mechanically-induced equilibrium, we also studied the kinetics and equilibrium properties of protein L in the presence of 3 µM DnaK and 5 mM ATP. Similar to DnaJ, DnaK has also been observed to increase the refolding MFPT (Fig. 2D), while significantly decreasing the unfolding MFPT than in its absence (Fig. 2E). Similar to DnaJ, the FP has also been observed to decrease from 0.5 to 0.2 in the presence of DnaK, suggesting a ∼56% reduction (Fig. 2F). However, this is less pronounced than in the presence of DnaJ, where FP is reduced by approximately 70%. We followed the same experimental approach to monitor the mechanical activity of SecB, a well-known chaperone to bind molten globule state and thus delay the formation of stable tertiary contacts^16, 50, 51^. We observed that very similar to DnaJ and DnaK, SecB hinders the protein folding by increasing the MFPT refolding, while decreasing the MFPT unfolding (Fig. 2G and 2H), which is also reconciled from the downshift of half-point force during FP analysis with SecB (Fig. 2I).

### The Synergistic action of DnaKJE on folding of protein L

So far, the experiments performed showed that individually DnaJ and DnaK chaperones significantly decrease the refolding ability of the substrate protein L under force and act as holdase (Fig. 2). We, therefore, sought to understand the effect of the whole complex, DnaKJE (comprised of DnaK, DnaJ, GrpE, and ATP), on the folding landscape of protein L. To systematically explore that, we unfolded and refolded the protein L octamer in the presence of chaperone complex (1 µM DnaJ+3 µM DnaK+5 µM GrpE+5 mM ATP+10 mM MgCl_2_) and compared with the DnaJ activity and control (Fig. 3A-3C).

**Figure 3:**
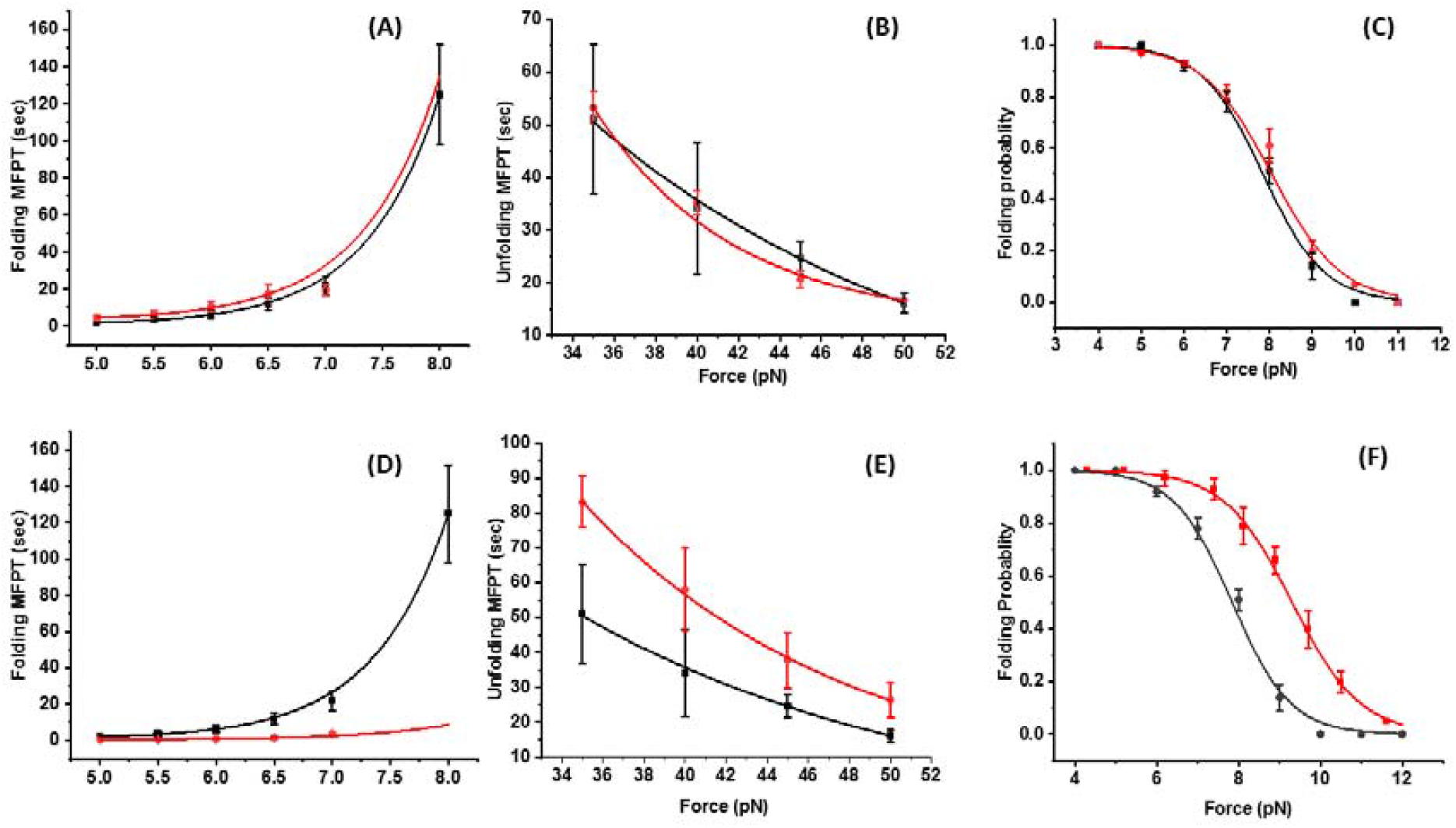
(A-C) DnaKJE chaperone complex shows a neutral effect on the folding ability in protein L: **(A) Differences in refolding MFPT:** MFPT of refolding are plotted as a function of force with DnaKJE chaperone complex (red) and in the absence of any chaperones (control, black). Protein L shows similar refolding time both in the presence of DnaKJE complex and control. Eight individual trajectories are measured and averaged. Error bars are s.e.m. **(B) Differences in unfolding MFPT:** Similar to refolding MFPT, unfolding MFPT is very similar to that of the control. More than five individual trajectories are measured and averaged. Error bars are s.e.m. **(C) Comparison of folding Probability:** Folding probabilities (FP) with DnaKJE chaperone system (red) and control (black) are plotted within 4-11 pN range, where FP with DnaKJE complex has been observed to overlap closely with the control. Data points are calculated using >2500s and over five molecules per force. Error bars are s.e.m. All the experiments are done in the presence of 5 mM ATP and 10 mM MgCl_2._ The buffer is changed in every 30 minutes to secure sufficient supply of ATP. **(D-F) DsbA dependency of folding properties of protein L: (D) MFPT of refolding:** DsbA positively modulates the refolding kinetics by speeding up the time to refold all the eight domains. MFPT refolding is much lower in the presence of DsbA (red) in comparison to control (black). Eight individual trajectories are measured and averaged. Error bars are standard error of the mean **(E) MFPT of unfolding:** DsbA slightly changes MFPT unfolding. Ten individual trajectories are measured and averaged. Error bars are standard error of the mean. **(F) Folding efficiency:** DsbA greatly increases the folding probability and the observed effect is most profound at 9 pN force. Folding probability of protein L in the presence of DsbA shown in red, whereas in absence of DsbA shown in black. Data points are computed using >2500s and over five molecules per force. Error bars represent s.e.m.

Notably, we found that in the presence of the DnaKJE complex (red), MFPT-refolding and unfolding closely overlap with that of control (Fig. 3A and 3B). Likewise, the folding probability of protein L in the presence of DnaKJE complex is comparable with that of control (Fig. 3C). In summary, these data nicely demonstrate that the whole DnaKJE complex simply allows protein L folding with the same efficiency, as in the absence of any chaperones, but better than individual DnaJ or DnaK. It is also important to note that, without GrpE, protein L in the presence of DnaKJ system (DnaJ/DnaK/ATP) is still working as a holdase (Supplementary Figure 6).

### Mechanical foldase activity of DsbA

The periplasmic oxidoreductase enzyme DsbA has a well characterized function to transfer the disulfide bond to substrate proteins by reducing its catalytic *CXXC* motif and helps in the folding of cysteine containing proteins^52–56^. Interestingly, few studies revealed the role of DsbA in transporting cysteine free proteins through translocon pore, suggesting its plausible chaperone activity independent of oxidoreductase activity^52, 57, 58^. Therefore, we explore the mechanical chaperone activity of DsbA by using our substrate protein L, a cysteine free globular protein. Unlike DnaJ or SecB, the refolding kinetics is greatly increased in the presence of DsbA, while slight change was observed in unfolding kinetics (Fig. 3D and 3E). At a given force, an efficient folding ability in proteins has been observed in the presence of DsbA (Fig. 3F) and the most profound effect is observed at 9 pN force. From the folding probability and MFPT calculations, we found that DsbA completely tilts the folding landscape towards the folded state, mostly by destabilizing the unfolded state. This finding is consistent with the accelerated protein folding process under force by DsbA assistance. The MFPT values has been cross-checked by ANOVA analysis and showed significant difference (Supplementary Figure 14).

The decrease in the unfolding kinetics indicates that DsbA is interacting with the folded/collapsed state of the substrate protein. This is exceptional for chaperones, however, common for thioredoxin proteins, including DsbA. Thioredoxin group containing proteins (for example PDI, Thioredoxin, DsbA) have been shown to interact with both the folded and unfolded states for disulfide bonded proteins^7, 45, 57^. After interacting with the unfolded state, it cleaves the disulfide bond, whereas it assists to reform the disulfide bond at low forces with the folded/collapsed state.

The equilibrium free energy of mechanical folding can be determined from the FP values by equation 1 and 2. These free energy values explain the mechanical foldase and unfoldase behavior of chaperones. For example, at 8 pN, the folding free energy difference of protein L is −0.04 kT (in the absence of any chaperones). This folding free energy has been observed to modulate upon the addition of chaperones. For example, it increases to +1.6 kT with DnaJ and +1.3 kT in the presence of DnaK, while decreases to −1.3 kT in the presence of foldase chaperone DsbA. These data reconcile that at force-induced equilibrium conditions, unfoldase chaperones compromise the refolding ability of substrate proteins by tilting the energy landscape towards the unfolded state, and in contrast the foldase chaperones shift the folding landscape towards the folded state. These free energy difference values of few kT seems to be small in comparison to the bulk experiments, however for single molecule experiment, few kT free energy difference are significant. For example, at equilibrium force condition of 8 pN with DnaK, the folding probability is decreased from 0.51 to 0.2, while in the presence of DsbA, it increases to 0.8. This implies that the population of folded state increased by 60% by foldase chaperone, whereas for the unfoldase chaperone, the population of folded state decreased by 60%. Our data clearly shows that folding probability values, associated to these substantial changes in folding efficiency are certainly arising from the chaperone effect and not from any thermal fluctuation, which are clearly observed from the folding probability.

### Mechanical chaperone activity of thioredoxin domain containing proteins

DsbA, possessing a *CXXC* motif (thioredoxin domain), shows a strong chaperone activity, which led us to further investigate whether this phenomenon is generalized to all thioredoxin domain containing proteins. We test two independent oxidoreductase proteins: full length PDI and Trx. Both of them are unable to show extra mechanical folding behavior as observed in the presence of DsbA (Fig. 4). This concludes that tunnel associated thioredoxin proteins like DsbA has a special mechanical activity by which they generate extra mechanical power to help the shipment of proteins from the cytoplasm to periplasmic side.

**Figure 4:**
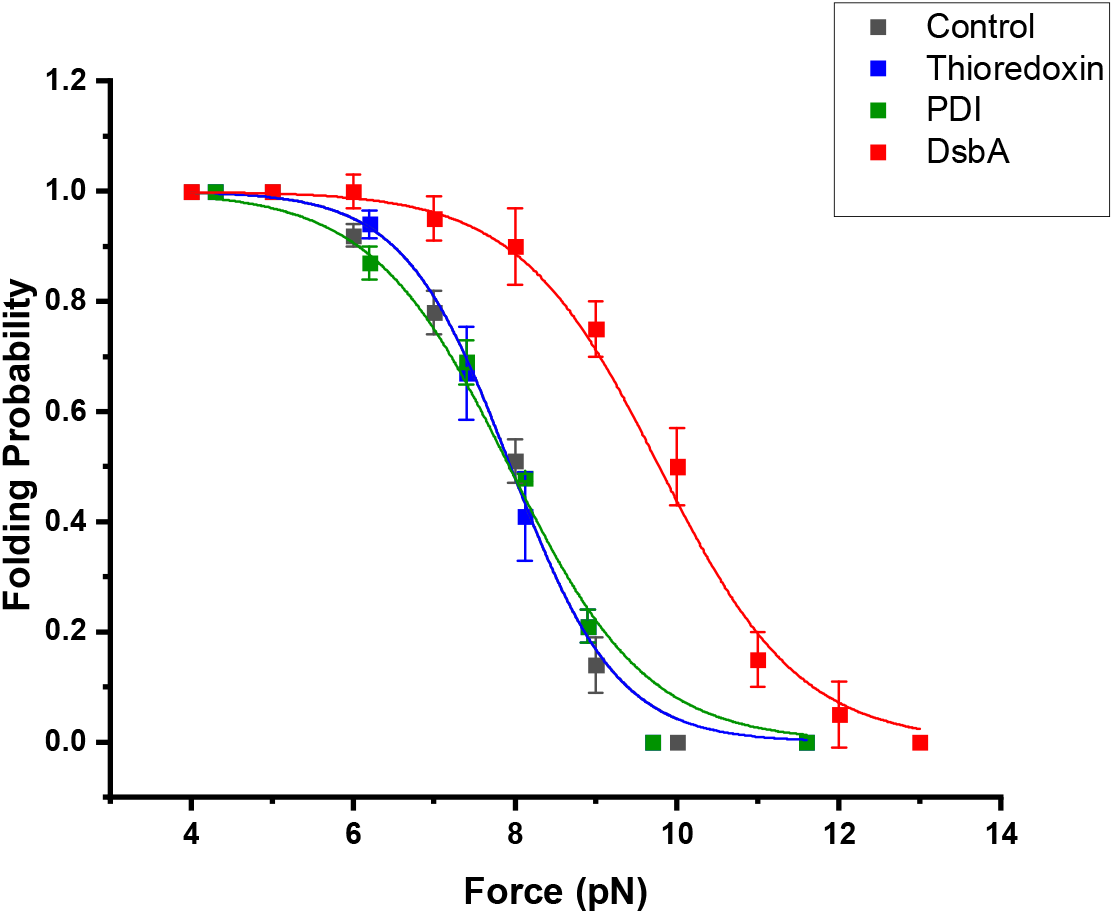
Comparison in folding probability of different thioredoxin domain containing proteins: Folding probability of protein L is plotted against force in the presence of full length PDI (green), Trx (blue) and DsbA (Red). Full length PDI and Trx, being cytosolic thioredoxin domain containing proteins, have only been observed to restore the folding efficiency in protein and overlaps with folding probability in control, whereas the tunnel associated counterpart DsbA not only restores but also accelerate the refolding yield in proteins under force. For example, at 8 pN, in the presence of both PDI and Trx, the folding probabilities are 0.5, respectively, while it increases to 0.8 in the presence of DsbA. Each data point is calculated using >2500s and over six molecules per force. Error bars represent s.e.m.

### Comparative analysis of chaperone activity on a single protein L

We represented a comparative analysis of how mechanical chaperones modulate the energy landscape and folding dynamics on a single protein L under equilibrium force condition at 8 pN (Fig. 5). In absence of chaperones, the polyprotein hops between its 4^th^ and 5^th^ folded state, generating a folding probability of approx. 0.5 (black, Fig. 5A). Further addition of mechanical chaperones shifts this folding transition either towards folded state or unfolded state. For example, in the presence of DnaJ the same polyprotein hops between its 1^st^ and 2^nd^ folded state (red trace, Fig. 5A) while addition of DsbA, increases this intrinsic folding ability to show a folding-unfolding transition between the 6^th^, 7^th^ and 8^th^ folded states, (blue, Fig. 5A). We compared the folding probability as a function of force with different chaperones as shown in Fig. 5B. In the presence of mechanical foldase chaperones such as, TF and DsbA, a rightward shift in folding probability is observed, representing a higher refolding yield in polyprotein whereas in the presence of other chaperones like DnaJ, DnaK and SecB individually, the half-point force is shifted towards the lower force regime, suggesting a compromised ability of proteins to remain folded under force. Indeed, these results collectively suggest the chaperones tilts the equilibrium folding landscape by modulating the intrinsic ability of protein folding under force.

**Figure 5:**
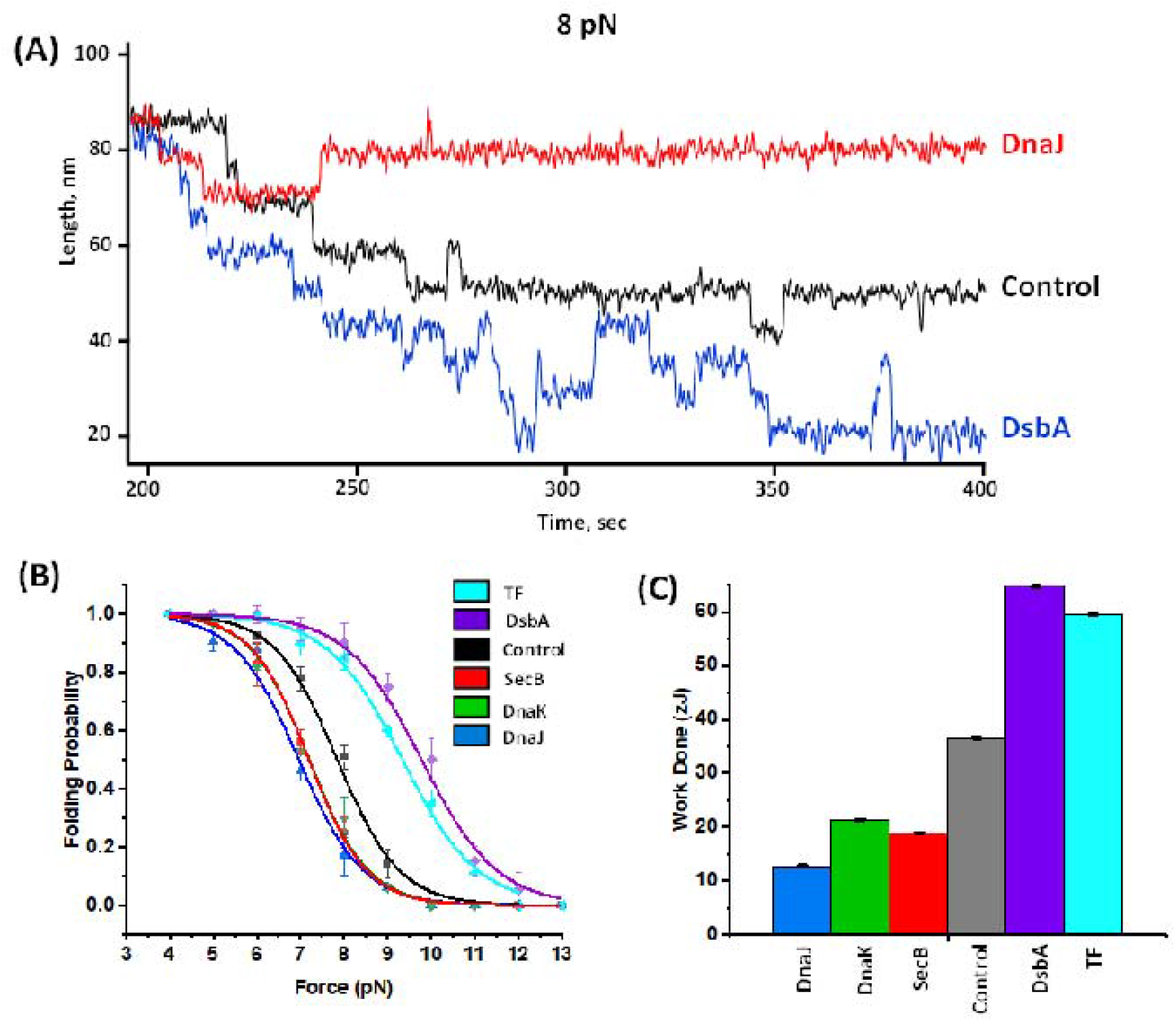
Mechanical activity of different chaperones at equilibrium force of 8 pN: **(A) Comparisons in folding transitions in the presence of different chaperones:** In absence of any chaperones (control, black trace), the polyprotein hops between its 4th and 5th folded states while introducing chaperones shift this folding transitions either towards folded state (downward) or unfolded (upward) state. For example, in the presence of DnaJ (red trace) protein L show a folding transition between its 1^st^ and 2^nd^ folded states while addition of DsbA (blue trace), increases this intrinsic folding ability, allowing a folding-unfolding transition between the 6^th^, 7^th^ and 8^th^ folded states. **(B) Variation in folding probability:** Folding probability is plotted against the force with different chaperones. In the presence of DsbA and TF, the folding probability is shifted rightward, suggesting a higher folding probability whereas the presence of chaperones such as, DnaJ, DnaK, and SecB retards the folding probability, representing a compromised refolding yield. The folding probability of TF is re-measured in the current buffer condition from Haldar et al. ^19^ Data points for each chaperone are calculated >2500s and over five molecules per force. Error bars represent s.e.m. **(C) Mechanical work output of different chaperones:** Mechanical work output is determined as the product of force, step size at that force and the folding probability. The calculated mechanical work for different chaperones is plotted where it has been observed that foldase like, TF and DsbA generate 59.5±0.1 and 64.8±0.1 zJ mechanical work where unfoldase such as, DnaJ, DnaK, and SecB generate 12.7±0.2, 21.4±0.1 and 18.9±0.1 zJ, respectively. Data bars are determined from the averaged FP and step sizes of more than eight molecules per chaperones at 8 pN force. Error bars represent s.e.m.

Protein folding against the force generates mechanical work^3, 4, 7^ for instance, titin protein produces 105 zJ of mechanical work during muscle contraction^3^. The work done by protein folding, under mechanical force, could be calculated by multiplying the force with step sizes as described previously by Rivas-Pardo et al^3^ and Eckels at al^7^. That means folding at the higher forces are larger and thus, can generate more work output. However, at higher forces, the folding probability is less. Therefore, a more useful measure for mechanical energy of protein folding is computed as a product of the work performed and the folding probability under a particular force. We observed that the chaperones maximally alter the mechanical energy in the equilibrium force range of 5-10 pN, by shifting the energy landscape either to the folded or unfolded state. Although chaperones could not influence the step size of protein L (Supplementary Figure 3), it largely affects the folding probability (Fig. 5B) and thus, invariably influence the mechanical work output. In case of mechanical foldase chaperones (like TF or DsbA), the work output is maximum while it is minimum for the unfoldase chaperones (like DnaJ, DnaK or SecB). The comparison of mechanical work output of different chaperones can be calculated for any force range, however pronounced difference could be achieved from the 7 and 8 pN force (Fig. 5 and Supplementary Figure 4). The mechanical work done values has been cross-checked by ANOVA analysis and showed significant difference (Supplementary Figure 14).

### Mechanical chaperone activity tested on Talin protein

To generalize our conclusion beyond protein L, we further extended our studies to a larger cytoplasmic protein talin R3-IVVI domain (124 residues), which is a well-characterized force-sensitive protein and has been reported to be higher mechanically stable than wild type R3^59, 60^. Furthermore, the structure of talin is completely different than that of proteinL: Talin is α-helix protein whereas protein L contains majorly β sheets and β-hairpins.

To monitor the mechanical activity of different chaperones, at first, we characterized the folding dynamics of talin at constant forces. Fig. 6 demonstrates the representative trajectories of talin folding dynamics, which is observed as extension changes of ∼20 nm and is strongly force-dependent. For example, talin domain mostly stays in the folded and unfolded state at 8.5 and 11 pN, respectively. At 10 pN, the domain occupies both the folded and unfolded states at almost equal populations. By applying constant force, we systematically investigated the mechanical effect of different chaperones on the folding dynamics of talin and observed very similar result to that with protein L. In the presence of unfoldase (DnaJ/DnaK), the folding dynamics shifts towards the lower force range: staying mostly in the folded state at 6.5 pN, unfolded state at 9.5 pN and populates both the states at 8 pN (Supplementary Figure 7A and 7B). In contrast, it shifts to higher force range in the presence of foldase chaperones. Such as with DsbA, the domain mostly populates in folded state at 13.5 pN, unfolded state at 16 pN, and at 14.5 pN, it occupies both the folded and unfolded states (Supplementary Figure 9A). Similar result is also observed with another foldase chaperone Trigger Factor (Supplementary Figure 9B). However, mechanically neutral chaperones such as, DnaKJE complex or full-length PDI have no additional effect on folding dynamics and exhibited same effect of promoting native dynamics in talin, as in the case of protein L (Supplementary Figure 8A and 8B).

**Figure 6:**
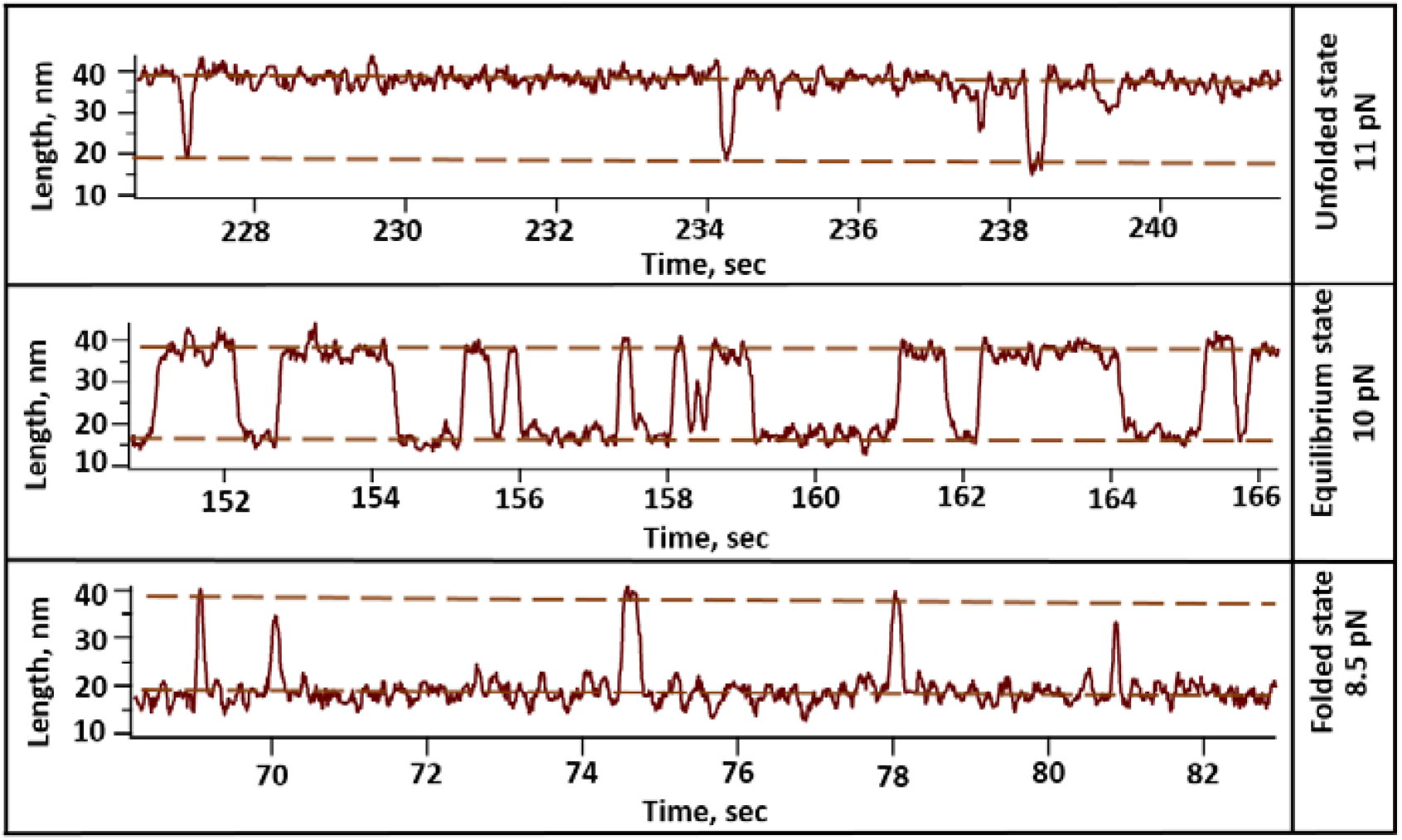
Representative trace of talin: Folding dynamics of talin domain is strongly force-dependent where the domain mostly remains unfolded at 11 pN, folded at 8.5 pN, and at 10 pN, both folded and unfolded states are equally populated.

The folding probability of talin, as similar to protein L, has also been observed to change in the presence of different chaperones. FP shifts to lower force range with unfoldase chaperones such as DnaJ, DnaK and DnaKJ complex, suggesting compromised refolding yields in talin (Fig. 7A, 7B and 7C), while in the presence of DsbA and TF, the FP shifts rightward, indicating higher folding ability in talin with the foldase chaperones, under force (Fig. 7F and 7G). However, FP of talin in the presence of mechanically neutral chaperones like DnaKJE complex and PDI, overlap closely with that in the absence of any chaperones or control (Fig. 7D and 7E). Due to the pronounced effect of chaperone on the force-dependent talin dynamics, their force regimes have been observed to alter significantly in the presence of different chaperones, which drastically affects the half-point forces. For example, in control (in the absence of any chaperones), the half-point is 9.8 pN, which decreases to 7.9 pN with DnaJ and increases to 14.9 pN in the presence of DsbA (Fig. 8A). Alterations in folding probability by chaperone interactions attributes to the underlying landscape mechanism that chaperones, similar to protein L substrate, reshapes the energy landscape of talin-unfoldase chaperones tilts the landscape towards the unfolded state which in turn reduces the folding probability in talin; whereas foldase chaperones shift the landscape towards the folded state by stabilizing it and thus, folding probability is increased under tension. Finally, we calculated the mechanical energy for the talin folding, in the presence of different chaperones. However, owing to distinct force regimes of talin dynamics with different chaperones, we calculated the mechanical energy as a product of constant FP=0.5, half-point force and the step size at that force. It has been observed that unfoldase decreases the mechanical energy by hindering the folding process under force, while foldases increase the mechanical energy by accelerating the folding process. For example, DsbA accelerates the folding process by increasing the refolding rate of talin, allowing the protein folding to generate ∼1.9 times more mechanical energy than in the presence of DnaJ (Fig. 8B). These results generalize our hypothesis of mechanical chaperone activity in modulating the folding process of mechanically stretched proteins.

**Figure 7:**
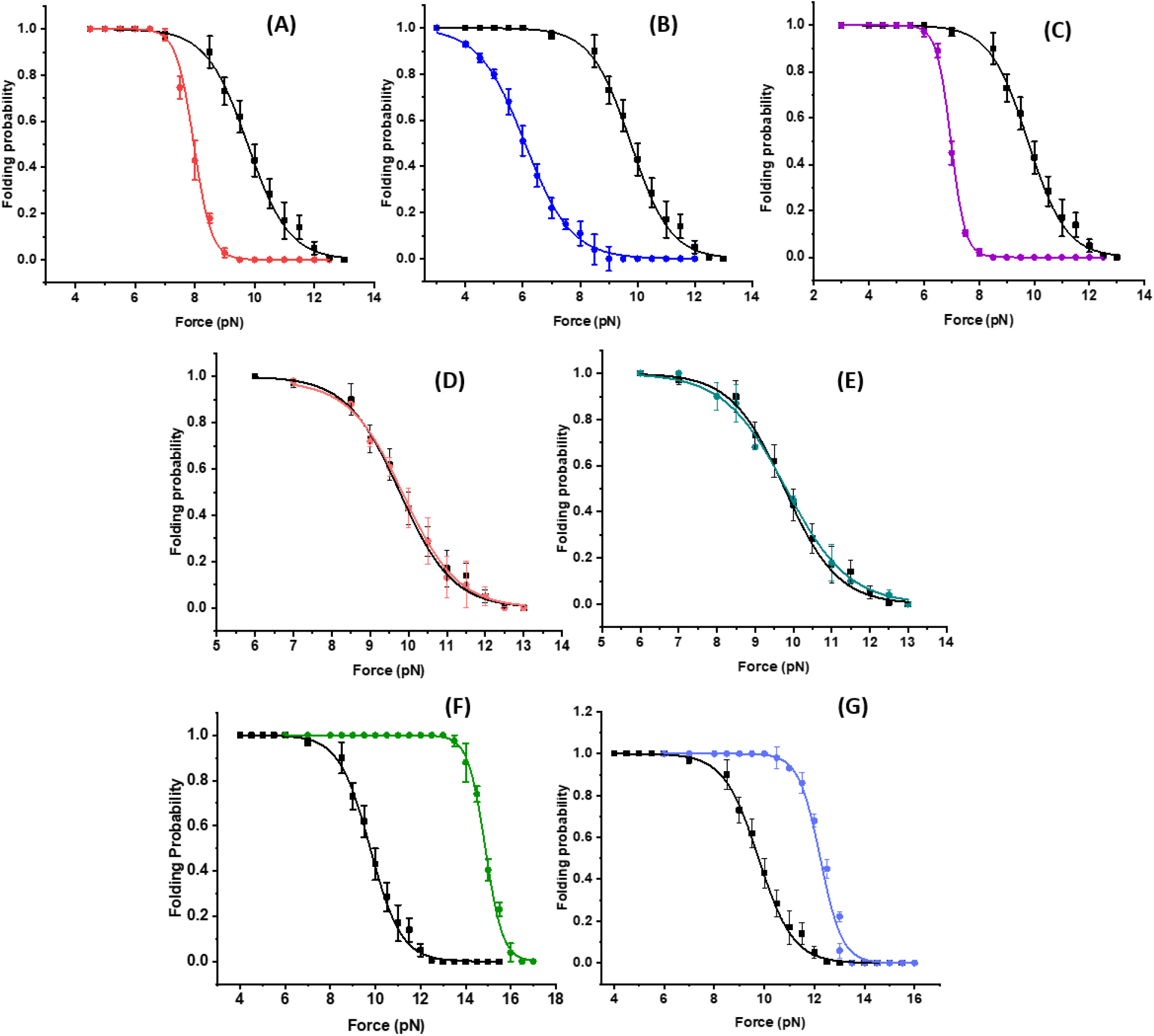
Chaperone-modulated folding probability of talin: **(A) Effect of DnaJ on the folding probability:** DnaJ shifts the folding probability of talin towards the lower force regime and the half-point force has also been observed to shift from 9.8 to 7.9 pN. Data points are calculated by using more than six individual protein molecules. Error bars are standard error by mean (s.e.m.). **(B) Effect of DnaK:** Folding probability of talin decreases significantly in the presence of DnaK and the half point force shifts from 9.8 to 6.1 pN. Data points are calculated by using more than eight individual protein molecules. Error bars are s.e.m. **(c) DnaKJ complex decreases folding probability:** In the presence of DnaKJ complex and ATP, the half-point force decreases to 6.9 pN. Data points are calculated by using more than six individual protein molecules. Error bars are s.e.m. **(D) DnaKJE complex promotes intrinsic folding ability:** In the presence of GrpE to DnaKJ complex and ATP, the half-point force reverts to 9.8 pN. Experiments with DnaKJ and DnaKJE complex were performed in the presence of 10 mM ATP and 10 mM MgCl_2_ buffer and the buffer is changed after every 30 minutes to keep the sufficient ATP supply. Data points ae calculated by using more than seven individual protein molecules. Error bars are s.e.m. **(E) Folding probability in the presence of PDI:** In the presence of PDI, the half-point force of talin remains 9.8 pN. Data points are calculated by using six individual protein molecules. Error bars are standard error by mean. **(F) DsbA increases folding probability:** DsbA significantly increases the folding probability by shifting the half-point force from 9.8 to 14.8 pN. Data points are calculated using more than six molecules per force. Errors bars are represented as s.e.m. **(G) Effect of TF on folding probability:** TF also shifts the folding probability of talin towards higher force regime and the half point force increases up to 12.2 pN. Data points are calculated by using more than six individual protein molecules. Error bars are s.e.m.

**Figure 8:**
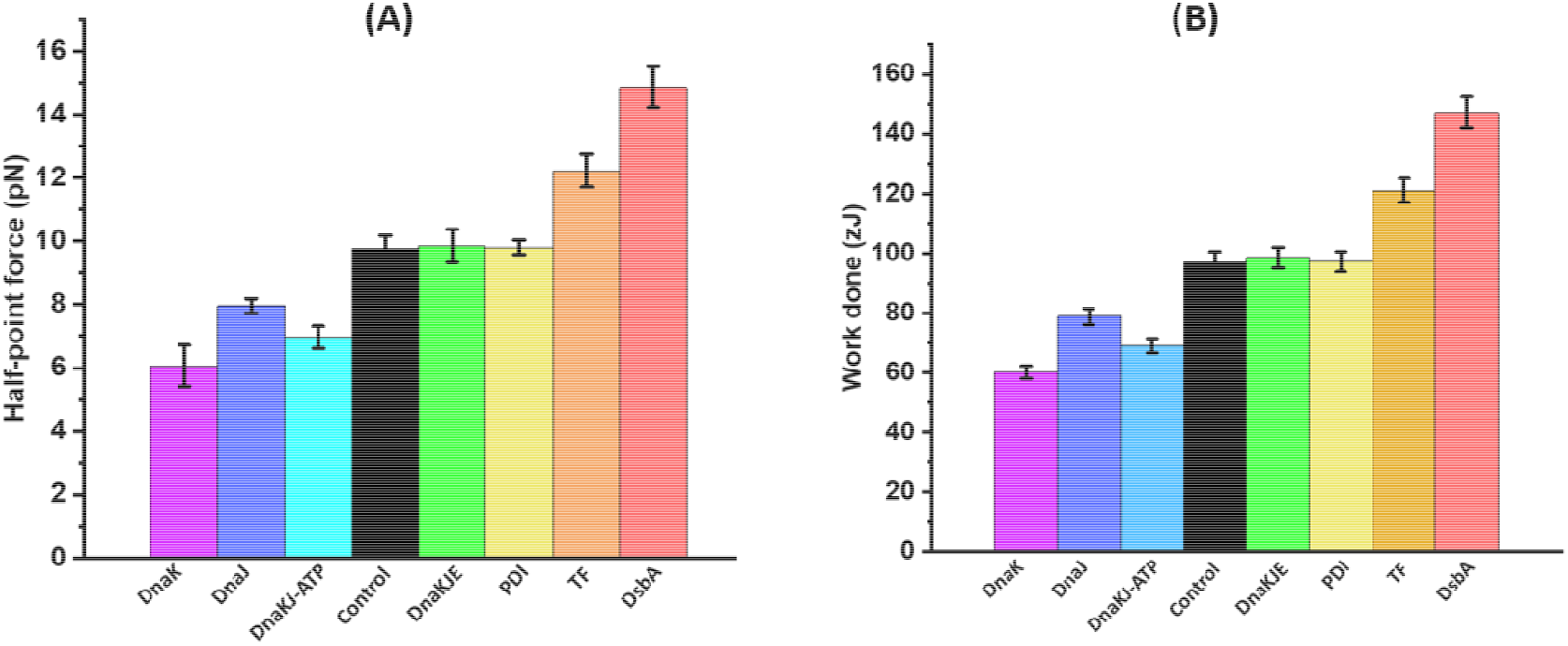
Mechanical work output of different chaperones at half-point forces: **(A)Half-point forces:** Due to the pronounced effect of chaperone on the force-dependent talin dynamics, their force regimes have been observed to alter significantly in the presence of different chaperones, which drastically affects the half-point forces. For example, in control (in the absence of any chaperones), the half-point is 9.8 pN, which decreases to 7.9 pN with DnaJ and increases to 14.9 pN in the presence of DsbA. **(B) Mechanical work output:** Mechanical work output is calculated as the product of folding probability, force and the step size at that force. Mechanical work done has been observed to decrease with the unfoldase chaperone, while increases with foldase chaperones. For example, in the presence of unfoldase chaperone like, DnaJ, DnaK and DnaKJ complex, the calculated mechanical work done is 78.8±2.7, 60±2.1 and 69±2.4 zJ, respectively; whereas in the presence of foldase such as DsbA and TF, mechanical work done is 147.2±5.2 and 121.±3.2 zJ, respectively. However, mechanically neutral chaperones such as DnaKJE complex and PDI have no additional effect on the folding process and thus generate similar amount of mechanical energy as that of control (98.6±3.4 for DnaKJE and 97.2±3.4 zJ for PDI). Error bars are s.e.m.

We have investigated the mechanical chaperone activity of eight different chaperones on two structurally distinct substrates i.e., protein L which predominantly contains β sheets and β-hairpins in their structure and talin domain containing four α helices packed against a buried hydrophobic threonine belt. Due to structural differences, these proteins exhibit different folding-unfolding pattern with distinct force-response, however, their folding landscape are modulated exactly similar during the chaperone interactions-unfoldase shifts the dynamics towards the unfolded state, while the foldase chaperones favor the folded state. Recently, different groups have also tested the unfoldase chaperone activity (DnaK/DnaJ) under force with immunoglobulin substrates, including titin I_27_ and Z1 proteins and observed similar results^34, 61^, which certainly coincides with our observation. Therefore, our results, along with other similar force-spectroscopy studies with structurally diverse substrates, have evidently strengthen and generalized our hypothesis of mechanical activity of chaperones.

## Discussion

Protein folding under force is a significant source of delivering the mechanical power to carry out different cellular processes^3, 4, 7, 62–64^. Although the chaperones are of central importance in many protein folding related phenomena, how they function and interact with the substrate protein under force remains poorly understood. Our results with the model substrate protein L and talin show empiric evidence and measurement that chaperones are able to generate mechanical work while assisting or hindering protein folding under mechanical loads.

Our single molecule experiments using constant force, provided a force-induced equilibrium condition, which revealed an additional question about the thermodynamics of mechanical folding pathways: How are the chaperones able to reshape the energy landscape? A possible explanation is protein shows intrinsic folding-unfolding transitions under equilibrium, where force acting as denaturant tilts the equilibrium free energy landscape towards the unfolded state. Chaperones, known to assist protein folding in multiple ways, most likely reshapes the mechanical folding dynamics by shifting the transition either towards folded or unfolded state. Therefore, we used two structurally distinct proteins-small globular protein L that mostly contains β sheets and β-hairpins in their structure and relatively larger talin R3-IVVI domain, containing four α helices. Despite of these structural differences, they undergo very similar folding landscape changes in the presence of chaperones.^19, 22, 37, 65, 66^ Unfoldase chaperones such as, DnaJ, DnaK and SecB have been observed to largely stabilize the unfolded state and reduces the folding probability. In contrast foldase chaperones, TF and DsbA have been observed to favor the refolding transitions under tension and show a more negative free energy towards the folded states.

Our experimental finding provides a global mechanical activity of chaperones in diverse protein folding pathways: starting from emerging out of the ribosome to getting localized in its destined cellular region. Throughout this pathway, the nascent polypeptides interact with several chaperones, and among them, TF is the first one. It interacts with the polypeptides while experiencing a mechanical tension, due to the constrained entropy, within the ribosome tunnel^19^. The ability of TF to fold against the force generates mechanical power that pulls the polypeptides out of the ribosome^19^. Other than assisting in folding process, TF carries its folded substrate to the SecA translocon motor, attached to the SecYEG translocon pore^50, 67–69^. However, TF presented polypeptide is not translocated by the SecA until it is unfolded by SecB, the go-to chaperone for translocation of several periplasmic proteins^40, 50, 51, 67, 70^. It is already reported that SecB binds to molten globule like structure of MBP^16^, however, in equilibrium, we found that SecB under mechanical constraints, binds to unfolded state and reduces the protein’s ability of conducting mechanical work by half (Fig. 5C). Interestingly, higher expression of DnaJ or DnaK compensates the absence of SecB in translocation of proteins, identifying another pathway where these chaperones direct the specific client proteins to the SecA translocon motor^40^. In addition to their functional similarity with SecB, DnaK and DnaJ individually prevents their substrate from folding, though their substrate specificity differs. Such as DnaK majorly binds to the molten globule state of mechanically stable ubiquitin, while DnaJ binds to completely unfolded polypeptide chain^34^. However, both of them shift the energy landscape towards the unfolded state, which prevent the folding and allows the polyprotein to do minimum mechanical work (Fig. 5C). This makes sense as otherwise, if the protein gets folded by itself, its translocation through the SecYEG pore might get hampered due to the geometric constraints or excess energy might be needed by SecA or other unfoldase to disrupt the tertiary folded structure^5, 16, 69^. However, DnaJ and DnaK along with the GrpE make the substrate folded by simply restoring its innate ability to refold under force, which has been clearly observed at equilibrium force condition (Fig. 3A-3C). So, if the substrate is a cytosolic protein, it will be assisted by foldase like, TF and DnaKJE complex otherwise, periplasmic proteins will be assisted by unfoldase like SecB, DnaK or DnaJ till they interact with SecA motor. Once received by the SecA, it keeps translocating the substrate which either remains unfolded by themselves or kept unfolded by other unfoldase, through the SecYEG pore at 20 amino acids or > 125-250 amino acids per ATP hydrolysis cycle, according to the destination of the proteins^69, 71^. In this translocon pore, the unfolded polypeptide experiences the same force^72^ as they face in ribosomal tunnel, which makes them unable to fold till they get to the periplasmic side. In the periplasmic side of SecYEG, tunnel associated chaperone DsbA folds the polypeptide and fastens the pulling of protein out of the tunnel by increasing its refolding kinetics. This enables the protein folding to generate almost two times more mechanical work at than in absence of DsbA (Fig. 5C). This excess work done by the protein folding might accelerate the translocation through the SecYEG pore.

Protein-chaperone interactions have been well-characterized by force spectroscopy studies and are subjects of intense debate. Bechtluft et al. has shown that SecB prevents the stable tertiary contacts and interacts only to molten globule state in MBP tetramer, suppressing the folding process^16^. TF has also been observed to protect partially folded structure by preventing the long-range interactions that form stable misfolded structure^17^. These studies suggested that chaperones redirect the folding pathway of proteins by binding and protecting their different conformations. Interestingly, a recent optical tweezers study has shown that TF promotes efficient folding of multidomain protein not only by reducing the misfolding but also by blocking denaturation process by unfolded domains^73^. Additionally, Mashaghi et al. showed that DnaK chaperone complex as a whole increases the refolding ability of the MBP tetramer, while individually DnaK stabilizes a compact MBP formation along with binding and stabilizing its unfolded conformation^15^. Chaperone activity of DnaKJE has also been studied on mechanically unfolded I27 domains by single molecule AFM where the chaperone complex suppresses the misfolding of the substrate protein. This single molecule study has also revealed that the canonical holdase DnaJ could function as foldase, increasing the folding at the expense of misfolding^36^. Overall, these reports had vividly described how in different ways chaperones assist substrates in folding, mostly by preventing the misfolded state of substrate proteins. Notably, mechanics of protein folding by Rivas-Pardo et al., presented a different hypothesis that protein folding under force could generate mechanical work output^3^. Chaperone, mostly known for their roles in protein folding, must interact to their substrate proteins under force. Despite of strong association with protein folding under force, how chaperones could modulate the mechanical work during diverse folding process remains largely unknown. In the present work, we demonstrate chaperones could behave differently if the substrate protein is under mechanical tension. For an example, our previous study revealed that TF, known to possess holdase function in the absence of force, behaves as a strong foldase while the client protein is under mechanical force^19^. A pronounced difference is also observed for DnaKJE complex: DnaKJE complex function as a strong foldase in the absence of force^74^, does not assist the substrate folding under force (Fig. 3C). However, individually DnaJ, DnaK, and SecB, both in the absence and presence of force, function as unfoldase (Fig. 2)^36, 50, 74^. Interestingly, we observed significantly strong foldase activity of oxidoreductase enzyme DsbA under mechanical force, as compared to the other cytoplasmic oxidoreductases such as, PDI and Trx. This is fascinating given that in the absence of force, DsbA shows a weaker chaperone activity than PDI in decreasing aggregation of non-disulfide containing substrates^26^, while exhibits approximately two-folds higher foldase activity than PDI in the presence of force (Fig. 4). This unique mechanical behavior of TF and DsbA originates from their subcellular locations: ribosomal exit tunnel for TF and periplasmic side of SecYEG translocon pore for DsbA. Both of these tunnels share similar confined geometry and exert the similar force on the unfolded polypeptides while emerging out of the ribosome or during translocation^6, 19, 75–77^. Our observations are strongly suggestive that this phenomenon could reveal a plausible biological scenario: during translation and translocation, foldase chaperones (DsbA or TF) fold the protein against the force to facilitate these processes. In contrast, cytoplasmic chaperones (PDI, DnaKJE or Thioredoxin) do not need to function under force and thus, they do not show the foldase activity under force. Thus, both the foldase and cytoplasmic chaperones show completely different activity in the presence and absence of force. However, transferring chaperones (SecB, DnaK or DnaJ) transfer the protein and to prevent the formation of the misfolded state, they favour the unfolded state both in the absence and presence of force. This provides an emerging insight of mechanical roles of chaperones: they can generate or consume energy by shifting the folding dynamics of the client proteins towards folded or unfolded state; suggesting an evolutionary mechanism to minimize the energy consumption in various biological processes.

## Materials and Methods

### Purification and expression of protein L

Sub-cloning of protein L and talin construct were engineered using *Bam*HI, *Bgl*II and *Kpn*I restriction sites as described previously^22, 60^. Protein L polyprotein was constructed by consecutively attaching eight protein L B1 domains, whereas in talin construct, is the same one used previously by Tapia-Rojo et al.^59, 60^ Polyprotein constructs were transformed into *Escherichia coli* BL21 (DE3 Invitrogen) competent cells. The cells were then grown in luria broth with carbenicillin for selection. After reaching optical density of 0.6-0.8 at 600 nm, cultures were induced with 1mM Isopropyl β-D-thiogalactopyranoside (IPTG, Sigma-Aldrich) and incubated overnight at 25°C. The cells were pelleted at 8000 rpm for 15 minutes and resuspended in 50mM sodium phosphate buffer pH 7.4 with 10% glycerol and 300 mM NaCl. 10 µM Phenylmethylsulfonyl (PMSF) as protease inhibitor, was added subsequently followed by lysozyme for membrane disruption. The dissolved pellet was ice incubated for 20minutes and mixed slowly at 4°C. The solution was then treated with Triton-X 100 (Sigma-Aldrich), DNase, RNase (Invitrogen) and MgCl_2_ and mixed in rocking platform at 4°C. The cells were homogenized at 19 psi and the cell lysate was collected after centrifugation. The protein was purified using Ni^2+^-NTA affinity chromatography of AKTA^TM^ Pure (GE healthcare). 20 mM imidazole containing binding buffer was used for the proteins for attaching to the column which was later eluted by 250 mM imidazole containing buffer. Biotinylation of polyprotein was done by Avidity biotinylation kit which was purified with Superdex-200 HR gel filtration column (GE healthcare) with 150 mM NaCl containing buffer.

### Expression and Purification of chaperons (DnaK, DnaJ, SecB, GrpE and DsbA)

Transforming the chaperone constructs in *Escherichia coli* DE3 cells (Invitrogen) provided us selected colonies which were then grown in luria broth media with respective antibiotics (ampicillin and carbenicillin 50 µg/ml) at 37° C. Cells were induced with 1 mM Isopropyl β-D-thiogalactopyranoside (IPTG, Sigma-Aldrich) and incubated overnight at 25°C for harvesting. For the purification of DnaJ, SecB and GrpE chaperones, the cells were harvested by centrifugation and pellet was resuspended in phosphate buffer (pH 7.4) with 10% glycerol and 300 mM NaCl. The solution was then mixed at 4° C with PMSF and lysozyme in it. Triton-X, DNase, RNase, and MgCl_2_ were added subsequently and mixed too. The cells were then burst open at 19 psi using French press followed by collection of cell lysate and purification using Ni^2+^-NTA affinity column of AKTA^TM^ Pure (GE Healthcare). The proteins were purified using 20 mM imidazole containing binding buffer and 250 mM imidazole containing elution sodium phosphate buffer (pH 7.4). Following the purification, proteins were separated from salts and concentrated using Amicon Ultra 15 (Merck). Full length DnaK chaperone was purified by Ni^2+^-NTA column, pre-loaded with Mge1 protein, as described elsewhere^78^. In case of DsbA, after pelleting the cell at 8000 rpm, it was collected in a pH 8.0, 50 mM Tris buffer with 20% (w/v) sucrose and 1 mM EDTA. DNase, RNase and PMSF were also added to this mixture followed by ice incubation. This resuspended solution was centrifuged where the supernatant (S-1) was collected while the pellet was dissolved in 20 mM Tris solution. This mixture was again centrifuged and supernatant (S-2) was collected. The two-supernatant containing protein lysate (S-1 and S-2) was purified by 1 M NaCl containing Tris buffer using AKTA^TM^ Pure protein purification system with Hi-Trap Q-FF anion exchange column (GE healthcare). 0.3% H_2_O_2_ was added to the purified proteins. Further size exclusion purification was done with Superdex-200 HR column of GE healthcare, with a running buffer containing 150 mM NaCl.

### Preparation of glass slide and coverslips

Magnetic tweezers experiments were carried out in a fluid chamber by sandwiching a glass slide and a coverslip (top and bottom), separated with thin layer of parafilm, providing an inlet and outlet system. Initially, the glass slides were washed with 1.5% Hellmanex III solution (Sigma Aldrich) and washed with double distilled water. The slides were then dipped in a solution containing methanol (CH_3_OH) and hydrochloric acid (HCl) following which they were soaked with sulfuric acid (H_2_SO_4_) and then washed. The slides were then treated in boiling water and then dried. For activating the glass surface 1% (3-Aminopropyl) trimethoxysilane (sigma Aldrich, 281778) was dissolved in ethanol and was silanized for 15 minutes. To remove the unreacted silane the glasses were ethanol washed and the surfaces were then baked at 65°C for ∼ 2 hour. Top coverslips were cleaned with 1.5% Hellmanex III solution (Sigma Aldrich) for 15 minutes and washed with ethanol followed by drying it in the oven for 10 minutes. The fluid chamber was then made by sandwiching the slide and coverslip, which was then filled with glutaraldehyde (Sigma Aldrich, G7776) in PBS buffer. After an hour, the chamber was washed with non-magnetic beads (2.5-2.9 µm, Spherotech, AP-25-10) reference beads followed by Halo-Tag amine (O_4_) ligand (Promega, P6741). The chambers were then treated with a blocking buffer (20 mM Tris-HCl, pH 7.4, 150 mM NaCl, 2 mM MgCl_2_, 1% BSA, 0.03% NaN_3_) to avoid non-specific interaction and kept in room temperature for 5 hours.

### Real-time magnetic tweezers experiment

Real-time magnetic tweezers instrument was setup on an inverted microscope with an oil-immersion objective that remains attached to a nanofocusing piezo. The magnet position was controlled through the voice-coil, located just above the sample chamber. The sample chamber was viewed using a white LED and the diffraction pattern images were collected with ximea camera (MQ013MG-ON). Gaussian fits were applied to determine the absolute position and the required data, corresponding to each step, was calculated by length histograms. The changes in length between the centers of two histogram peaks denotes the step size. The force calibration has been shown in detail in two of our parallel manuscript. As these are not published yet, we are providing them in the supporting information (Supplementary Figure 10-12). The magnetic tweezers experiments were carried out with 5 nM of protein L in a buffer containing 20 mM sodium phosphate, 140 mM NaCl and 10 mM ascorbic acid at pH 7.2. Ascorbic acid has been used as antioxidants to prevent the oxidative damage of the protein L^71^. 1 μM DnaJ, 3 μM DnaK, 60 μM oxidized DsbA, 10 μM PDI, 10 μM Thioredoxin and 1 μM SecB was used for the chaperone experiments. For the experiments with DnaKJE chaperone complex, the buffer was supplemented with 5 mM ATP and 10 mM MgCl_2_. Our result shows folding of protein L is not affected by 10mM ascorbic acid (Supplementary Figure 5). After passing the protein L through the chamber, the octamer was attached with a paramagnetic bead (Dynabeads M-270) to which the force is applied with a pair of neodymium magnets. Experiments were started after z-stack of the library of both the paramagnetic and reference beads were acquired, and displacements of z position were determined using the image processing method, as described previously^22^. Gaussian fits were used to measure the absolute position and the data corresponding to each step has been used to obtain the length histograms. The distance between the centers of two adjacent peaks is defined as the step size^22^. We applied different forces to measure the step size, describing the unfolding and refolding of single protein domains. Different force was applied on protein L to which the effect of different chaperones was observed.

## Acknowledgement

We thank Ashoka University for support and funding. S.H. thanks DBT Ramalingaswami Fellowship and DST SERB Core Research Grant for funding. We thank Dr. Koyeli Mapa of Shiv Nadar University for kindly sharing with us the clones of SecB, DnaK, DnaJ, GrpE proteins. We thank Dr. Kausik Chakraborty (Institute of Genomics and Integrative Biology) and Prof LS Shashidhara (Ashoka University) for the critical analysis of this work.

## Supplementary Information

### Folding probability calculations

We described the dwell time analysis of folding probability of protein L (Fig. 1B). Calculation of the folding probability using the eq. 1 mentioned in the result section.

**Supplementary Table 1:** State occupation at 7 pN force of protein L B1 domain in the absence of any chaperones.

| **Force = 7 pN** |  |  |
| --- | --- | --- |
| **Control** |  |  |
| **State I** | **t** | **It/N** |
| 8 | 0.03 | 0.03 |
| 7 | 0.646 | 0.565 |
| 6 | 0.136 | 0.102 |
| 5 | 0.092 | 0.057 |
| 4 | 0.062 | 0.031 |
| 3 | 0.000 | 0.000 |
| 2 | 0.000 | 0.000 |
| 1 | 0.000 | 0.000 |
| 0 | 0.000 | 0.000 |
| **FP = 0.8** |  |  |

**Supplementary Figure 1:**
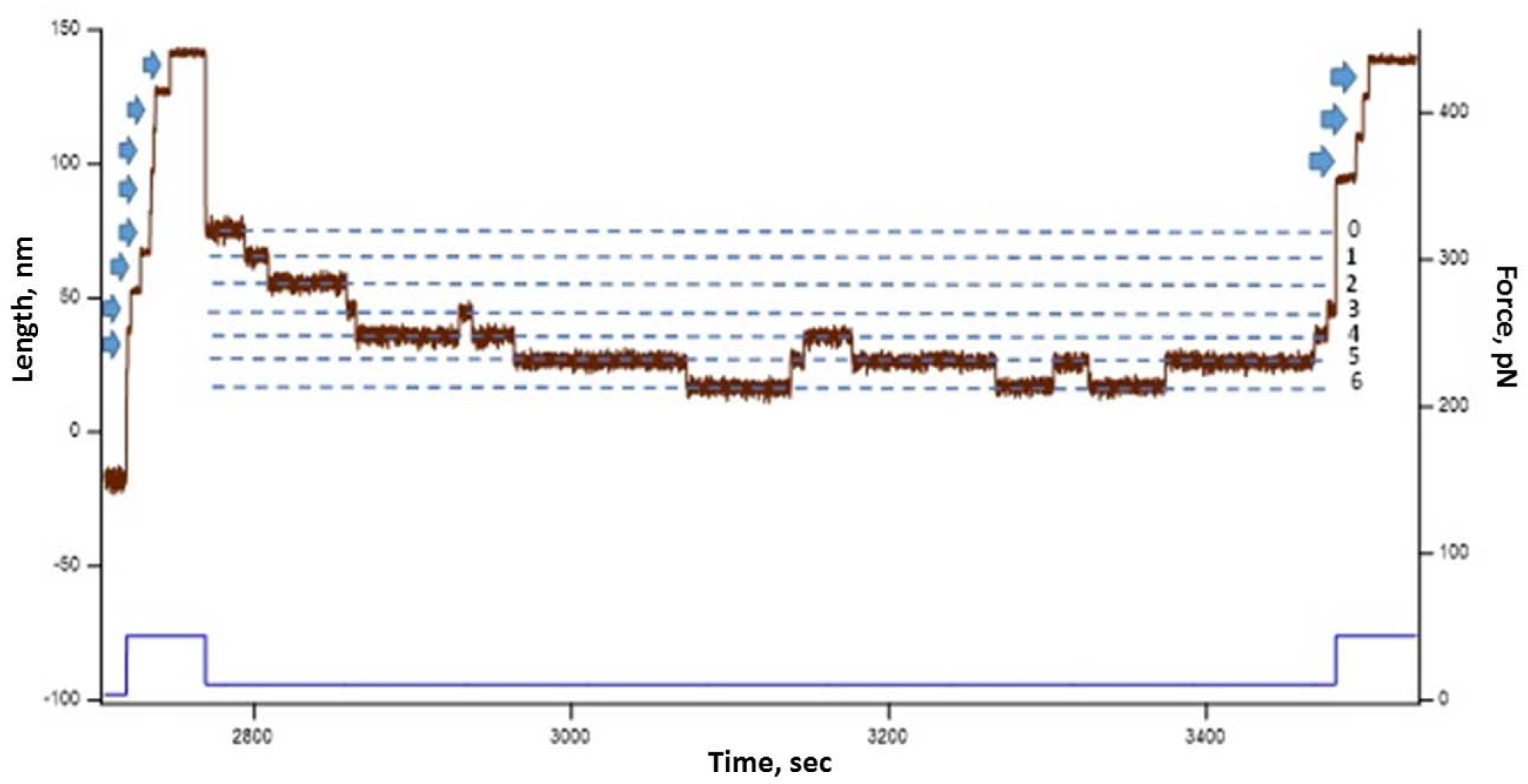
First, the polyprotein is unfolded at high force pulse of 45 pN, generating eight successive steps as single molecule fingerprint that followed by a refolding pulse at 11 pN. At 11pN, the domains show a continuous hopping between folded and unfolded state and then they are unfolded again at 45 pN. Before pulling at 45 pN, 3 domains folded at 11 pN and thus, when it is pulled at 45 pN, 3 distinct unfolding steps was observed.

**Supplementary Figure 2:**
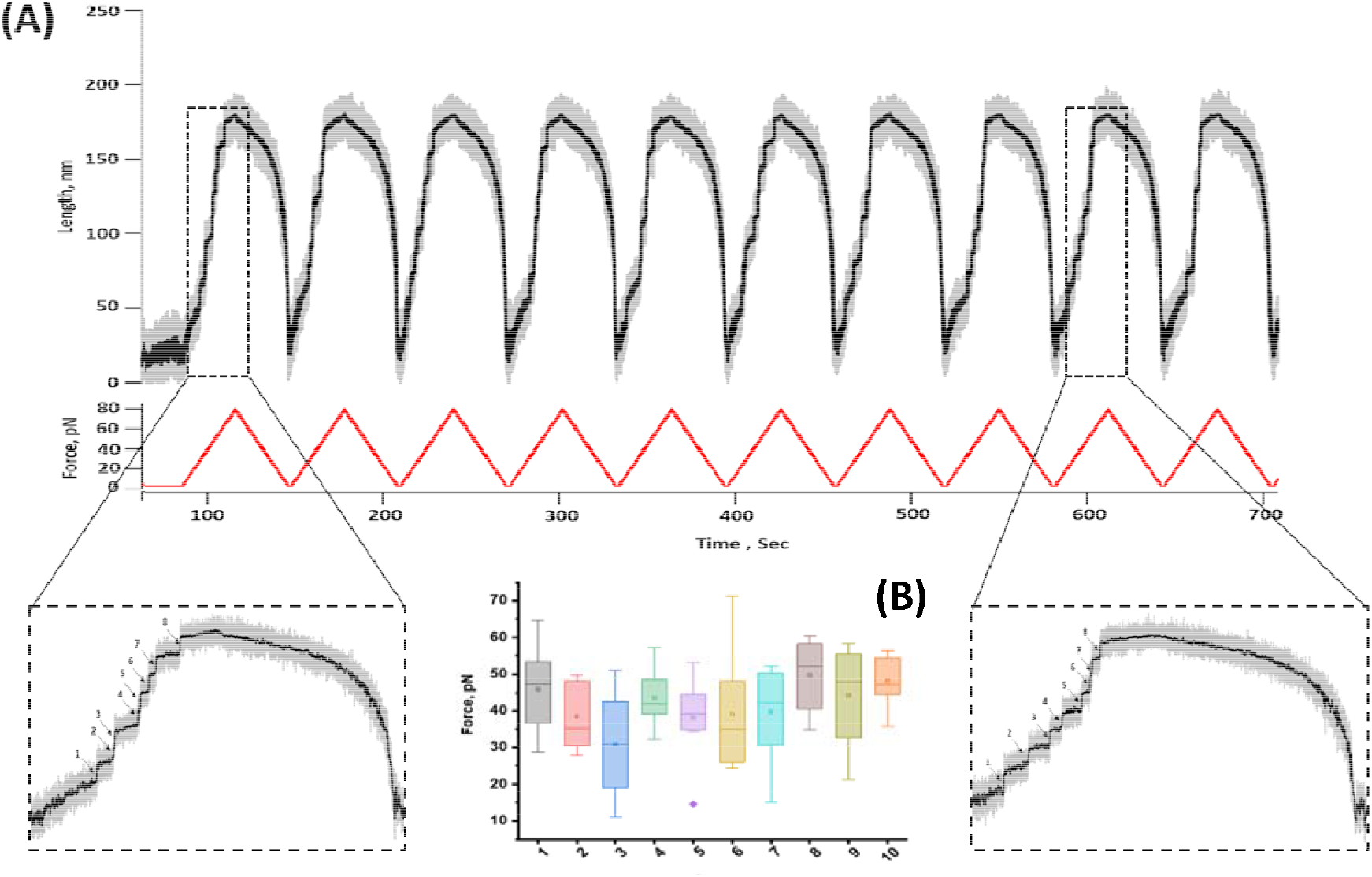
Mechanical strength of protein L octamer: **(A) Force ramp study:** Successive force ramp, from 4 to 80 pN at 2.5 pN/s ramp rate, are applied to detect the presence of any misfolded states. The unfolding traces are magnified to observe eight discrete step sizes throughout the successive ramp study. **(B)ANOVA analysis of average unfolding force of protein L with unfolding/refolding pulses:** Mechanical strength of polyprotein does not change with unfolding and refolding pulses, showing the absence of any misfolded state of protein L under force. ANOVA analysis shows that the differences between different population means are non-significant.

**Supplymentary Figure 3:**
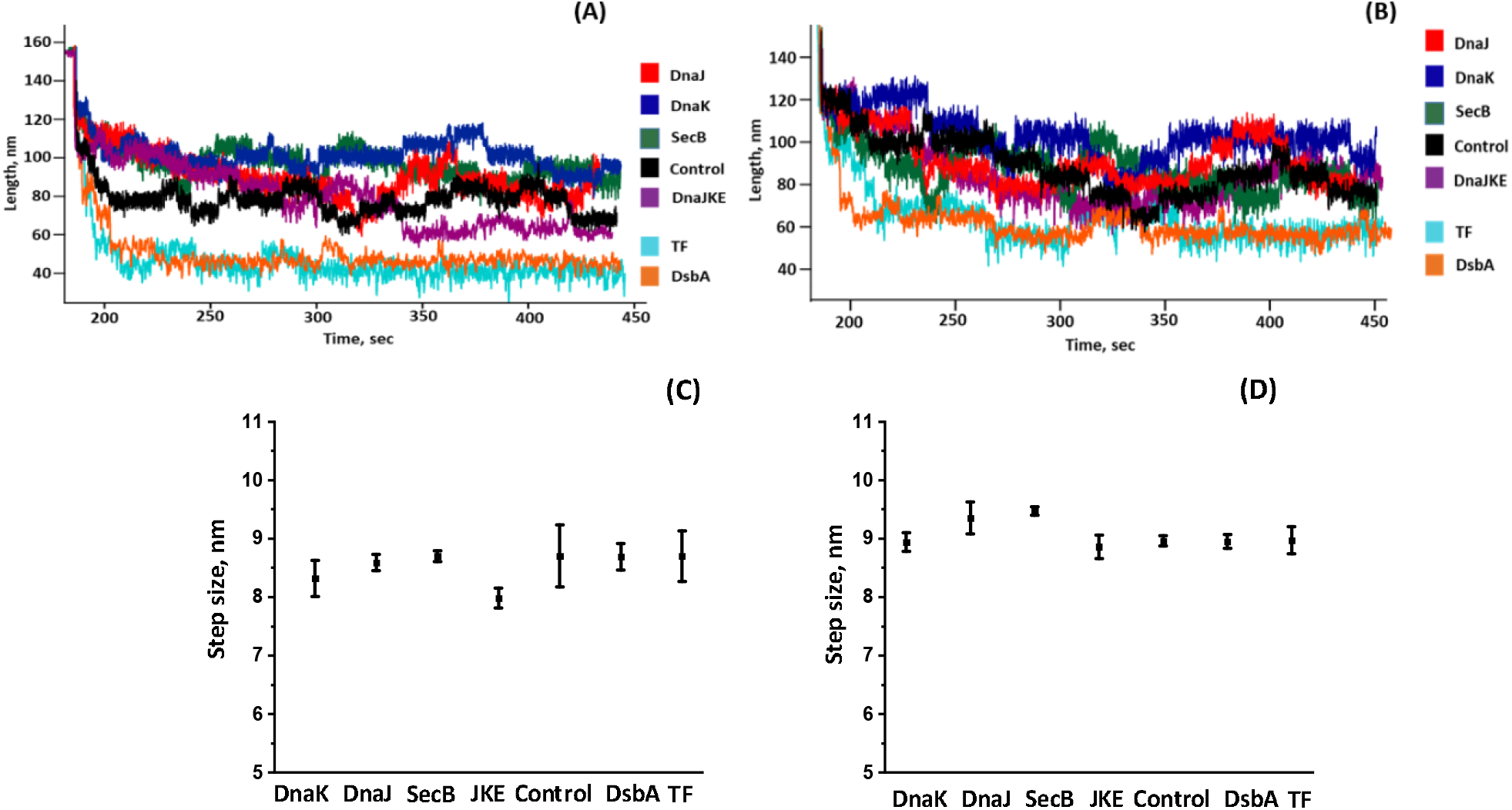
Protein L does not form any misfolded state: Protein L misfolding certainly affects their step sizes. To monitor that, equilibrium trajectories at 7 pN (A) and 8 pN (B) are analyzed critically analyzed in the presence and absence of any chaperones (control) to observe any changes in the step sizes, occurred due to misfolding events in protein L. We observe no significant changes in the average step sizes with different chaperones at both 7 pN (C) and 8 pN (D). Data points are determined from the step sizes of more than ten proteinL molecules per chaperones at 7 and 8 pN force. Error bars represent s.e.m.

**Supplementary Figure 4:**
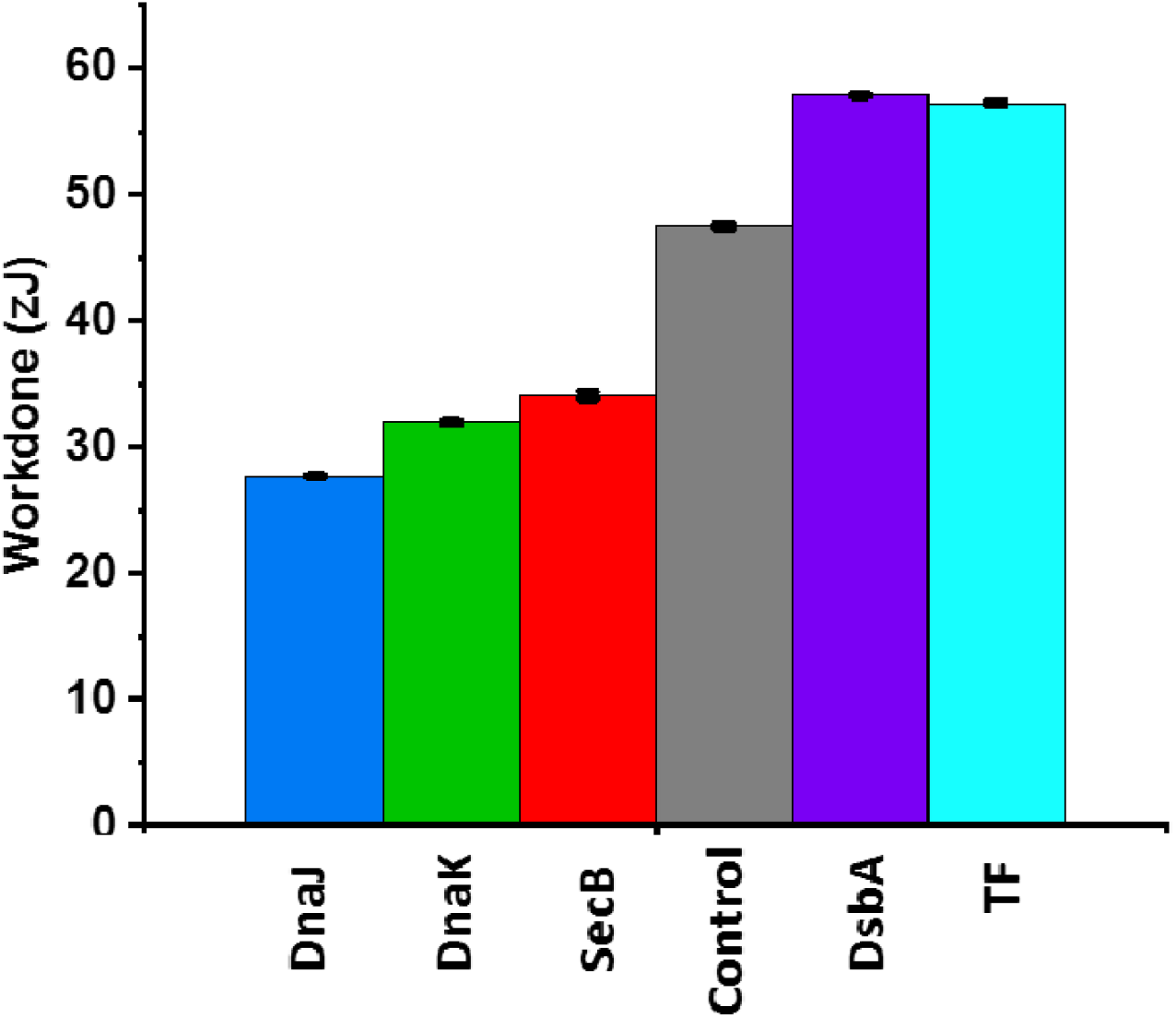
Mechanical work done by protein L folding at a force of 7 pN, in presence and absence of different chaperones: Work done has been measured for protein L in absence of any chaperones (Control, 47.5 zJ). While in presence of three different unfoldases DnaJ (27.7±0.1 zJ), DnaK (31.9 ±0.1zJ), SecB (34.1±0.3 zJ) and two foldases DsbA (57.2±0.1 zJ) and TF (57.2±0.1 zJ) at 7 pN force. Data bars are determined from the averaged FP and step sizes of more than five molecules per chaperones at 7 pN force. Error bars represent s.e.m.

**Supplementary Figure 5:**
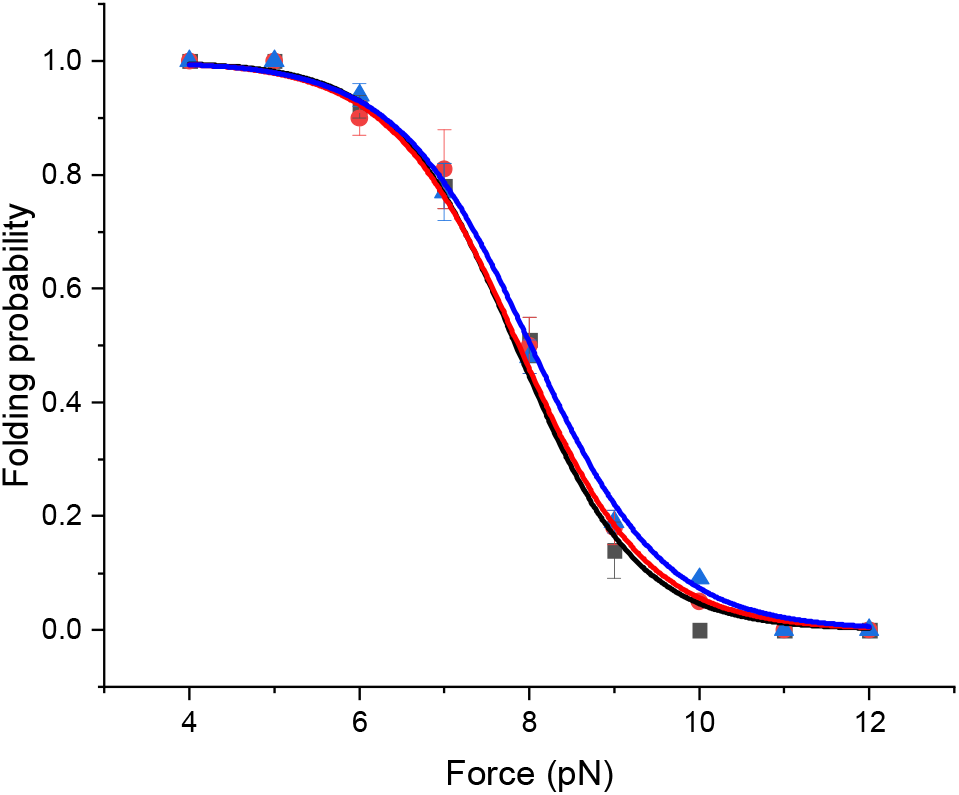
Folding probability of protein L in presence of 0 mM (red), 5 mM (black) and 10 mM (blue) ascorbic acid.

**Supplementary Figure 6:**
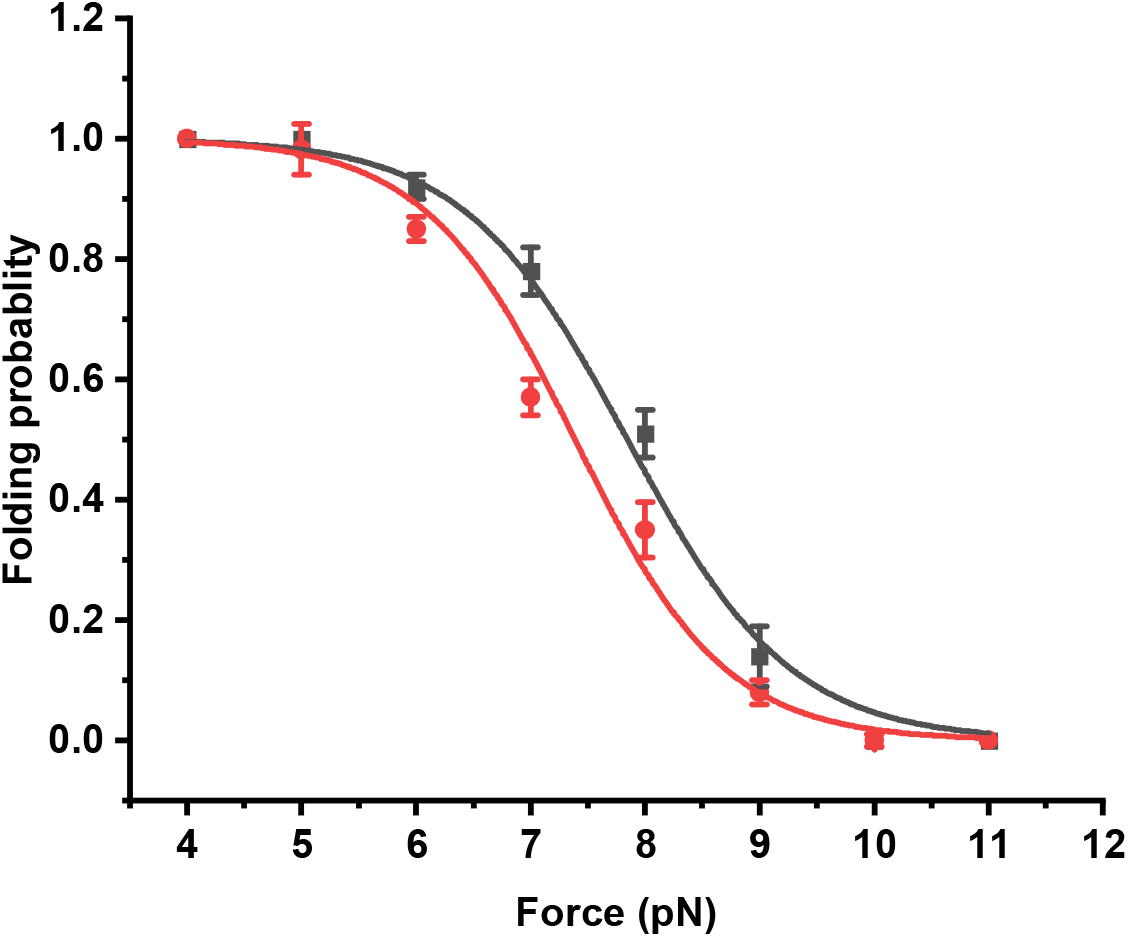
Folding probability of protein L in the absence (black) and presence of DnaK, DnaJ with 5 mM ATP and 10 mM MgCl_2_(Red). The buffer is changed every 30 minutes to secure sufficient supply of ATP.

**Supplementary Figure 7:**
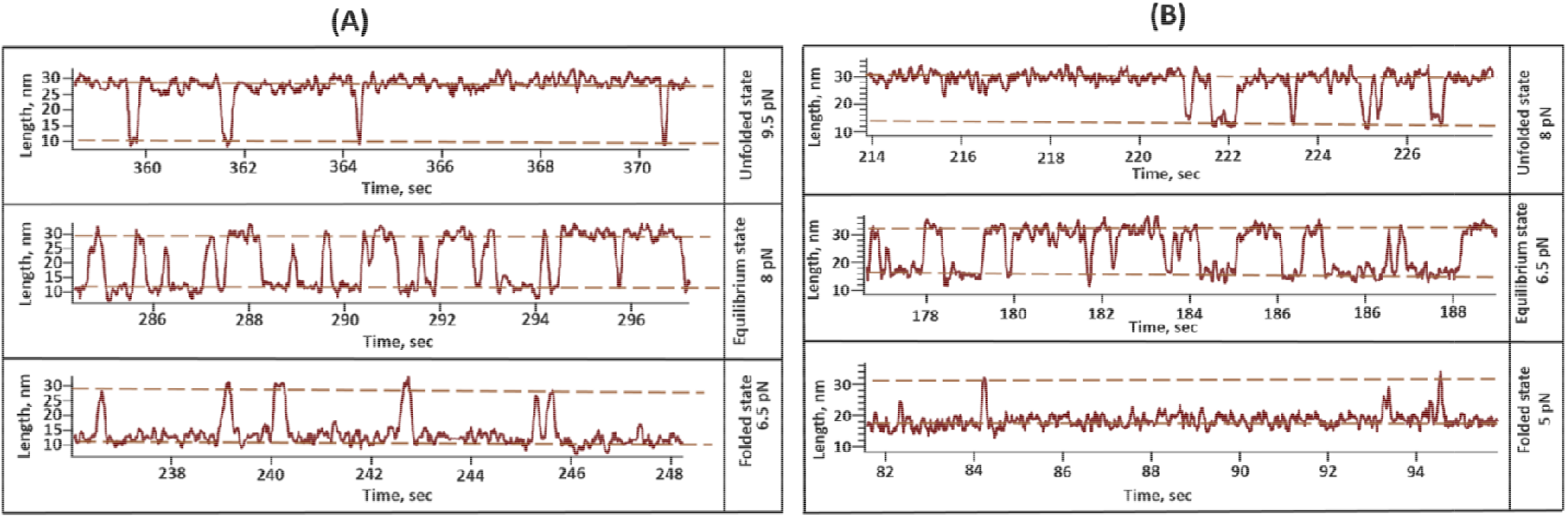
Unfoldase shifts the folding dynamics to lower force range: **(A) DnaJ:** Folding dynamics of talin has been shifted towards lower force regime in the presence of 1 µM DnaJ. At 9.5 pN it is mostly populated in the unfolded state and mostly folded at 6.5 pN. At 8 pN, it stays almost 50% in the folded state. **(B) DnaK:** The folding dynamics has also been observed to downshift with 3 µM DnaK. Talin remains mostly folded at 5 pN and unfolded at 8 pN and at 6.5 pN, it shows equal population of folding and unfolding.

**Supplementary Figure 8:**
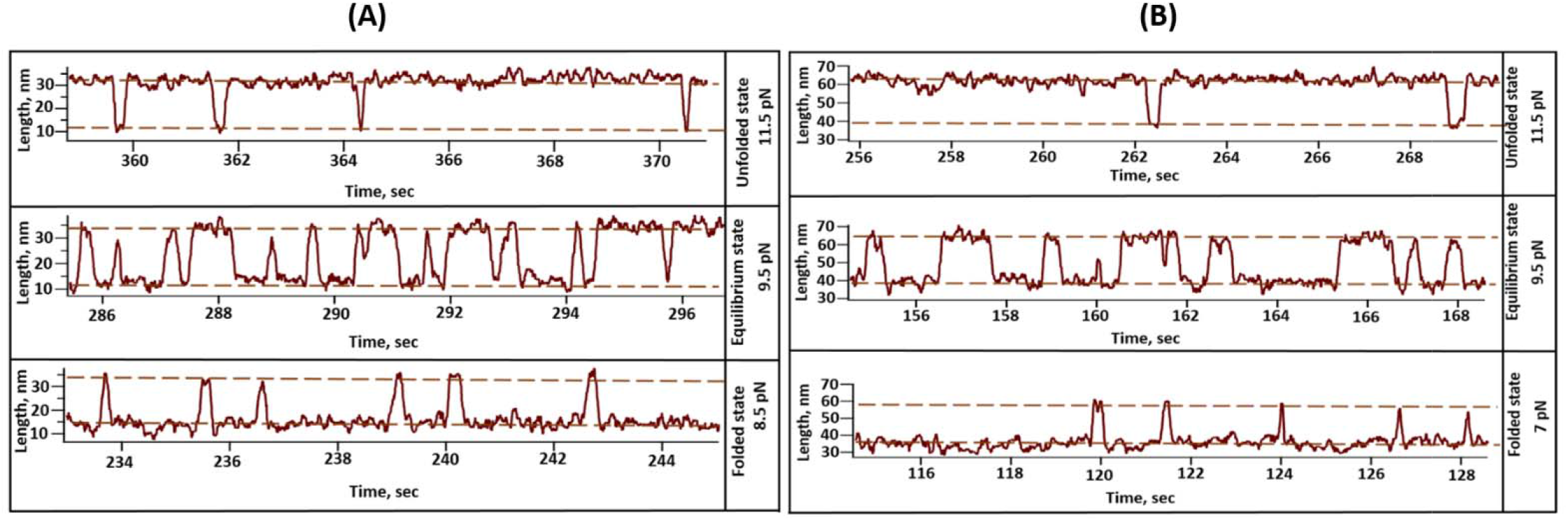
Mechanically neutral chaperones have no effect on the folding dynamics: **(A) DnaKJE complex:** The folding dynamics of talin is measured in the presence of 1 µM DnaJ, 3 µM DnaK, 5 µM GrpE, supplemented with 10mM ATP and 10 mM MgCl_2_. We observed that at 9.5 pN, the domain is equally populated in both the folded and unfolded states. **(B) PDI:** Similarly, for PDI, it has been observed that talin domain occupies both the states at 9.5 pN and thus, DnaKJE and PDI have no additional effect on folding dynamics and exhibited same effect of promoting native dynamics in talin.

**Supplementary Figure 9:**
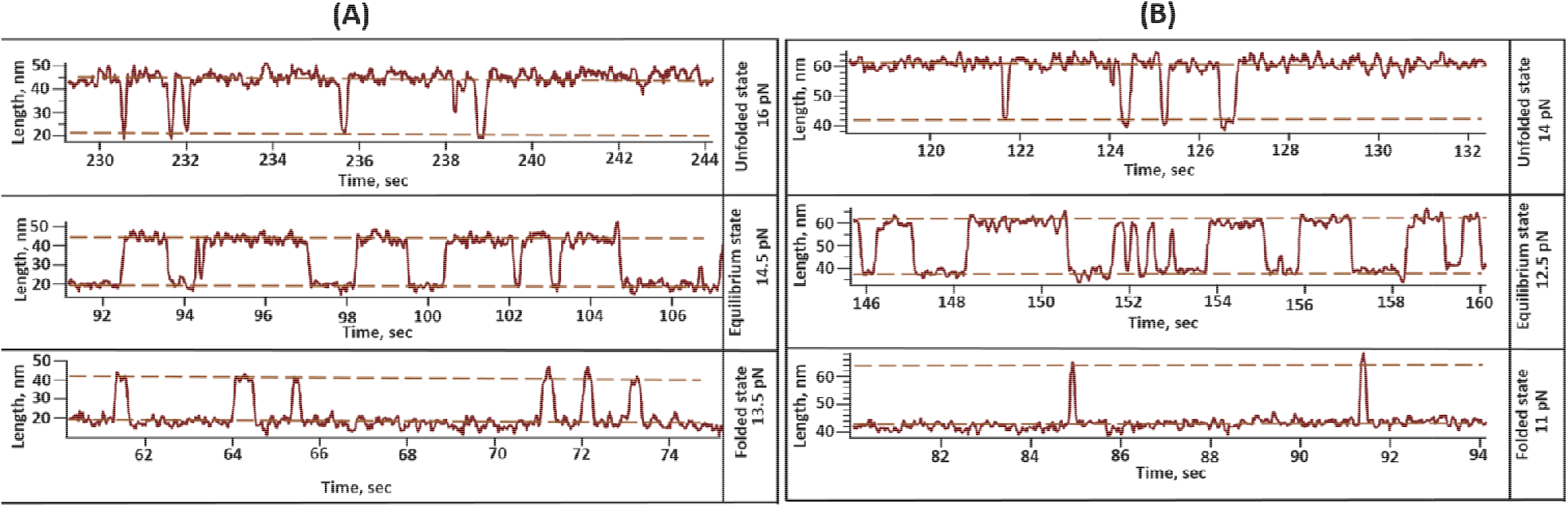
Mechanical foldase chaperones shifts the folding dynamics to higher force range: **(A) DsbA:** In the presence of 60 µM DsbA, talin mostly remains folded at 13.5 pN and unfolded at 16 pN and at 14 pN, it equally populated in folded and unfolded states. **(B) Effect of TF:** In the presence of TF, talin folding dynamics of talin has also been shifted towards higher force regime, where it shows mostly in the folded state at 11 pN and mostly in the unfolded state at 14 pN. It shows equally populated folded and unfolded state at 12.5 pN.

### Force calibration method in single molecule magnetic tweezers

In single-molecule magnetic tweezers, the force can be calibrated either by measuring the transverse fluctuation in case of large construct such as nucleic acids or by comparing protein unfolding extension to polymer elasticity models.^10^ As we have used small protein tether with step size with 6 to 20 nm, we pursued the calibration method, required for the protein construct.^1–9^ We confirmed our calibration method with conventional B-S transition of 565 bp dsDNA.^14^ We have measured the unfolding step sizes against the applied force in PBS buffer and fitted with the freely jointed chain model (Supplementary Figure 10) with *L_c_*=16.6±0.3 nm and *L_k_*=1.1±0.2 nm, which are consistent to the data from the previous publications.^4, 16^ Furthermore, to validate the force calibration accuracy, we fitted the applied force at different magnet distance to the magnet law (Supplementary Figure 11).^16^

Force calibration sensitivity has been monitored by imposing the deviation length values: 0.3 nm for *L_c_* and 0.2 for *L_k_* and observed that the effect of these deviations is within the 95% confidence level. Due to the bead size variation and alteration in attachment point to the paramagnetic beads, the applied force has approx. 10% uncertainty.^15, 16^ However, the M270 dynabeads have lower *cv* (coefficient of variation) of ∼2% than other beads, which could also confirm the minimal heterogeneity among M270 beads. It is imperative to adjust the voice coil in reproducible manner for the force application by the magnets. Here, we have used small globular protein as a tether and therefore, due to their higher corner frequency in y-fluctuation (or called transverse fluctuation), force calibration using thermal fluctuation can only be performed at lower force range at <15 pN, while the force calibrated by extrapolating the force calibration curve (Supplementary Figure 11).^15, 17^

### Force calibration validation

We cross-checked our calibration method from the folding probability data of protein L substrate and then compared our calibration with three different groups.^2–4^ The folding probability of protein L exhibits strong force-dependency and it has been monitored within 4 to 11 pN range. We observed that half-point force of protein L is ∼8 pN force and it perturbs significantly with 1 pN force deviation. For example, at 7 pN, the folding probability can increase by ∼56% (folding probability at 7 pN is 0.8), while decreases by ∼72% at 9 pN (folding probability at 9 pN is 0.1). Therefore, small alteration in applied force after calibrating, markedly perturb the folding efficiency of substrate protein under mechanical force. Here, we found protein L display similar half-point force at ∼8 pN, which strongly agrees with that reported in other studies and all the folding probability values are in well-agreement with these values (Supplementary Figure 12). Therefore, this accurate detection at sub-pN force range, which allow us to monitor the folding mechanics, could ensure its fiduciary measurement by our force calibration method.

**Supplementary Figure 10:**
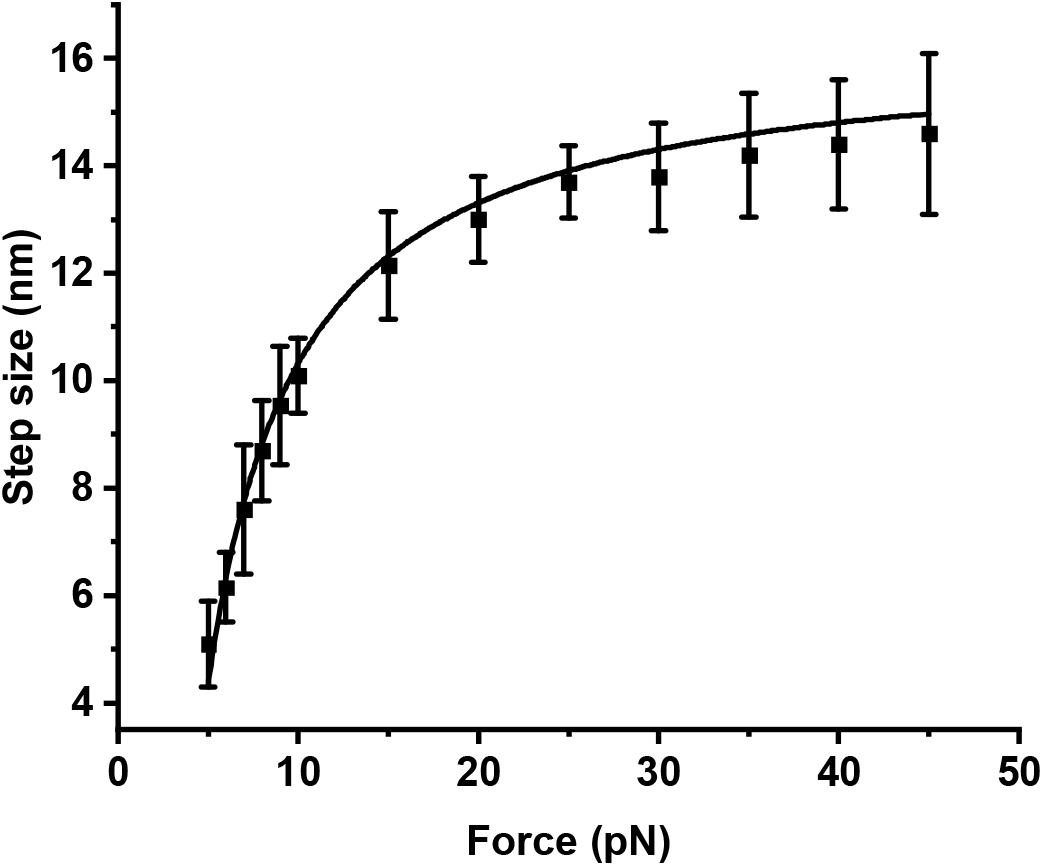
Step-sizes at different forces are fitted with freely jointed chain (FJC) model of polymer elasticity.

**Supplementary Figure 11:**
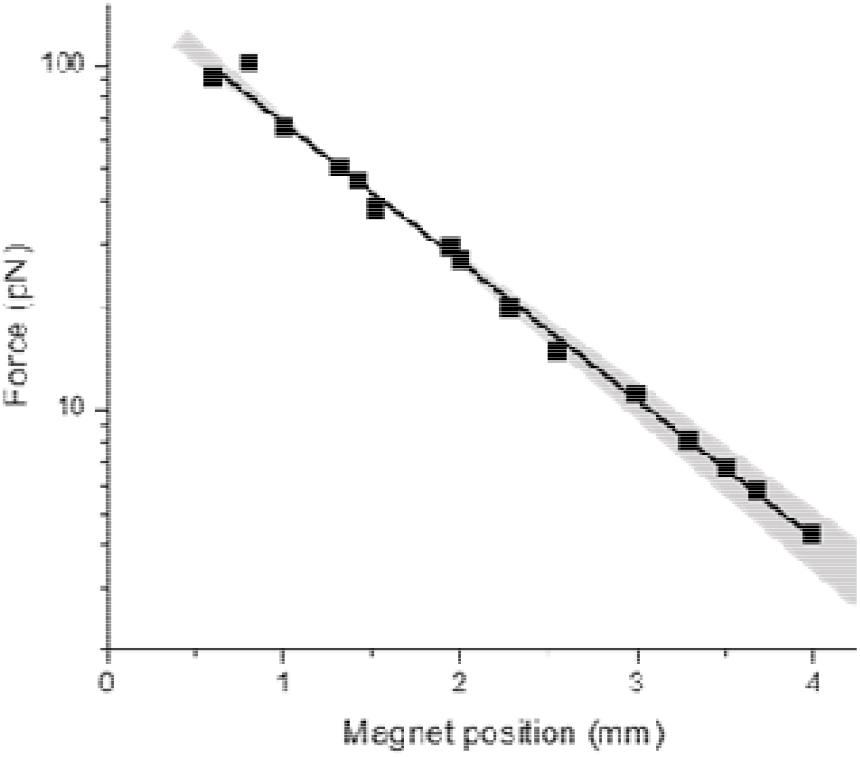
Force calibration by magnet law: We calibrated the force by the magnet law, which was proposed by Popa et al. (J. Am. Chem. Soc., 2016)

**Supplementary Figure 12:**
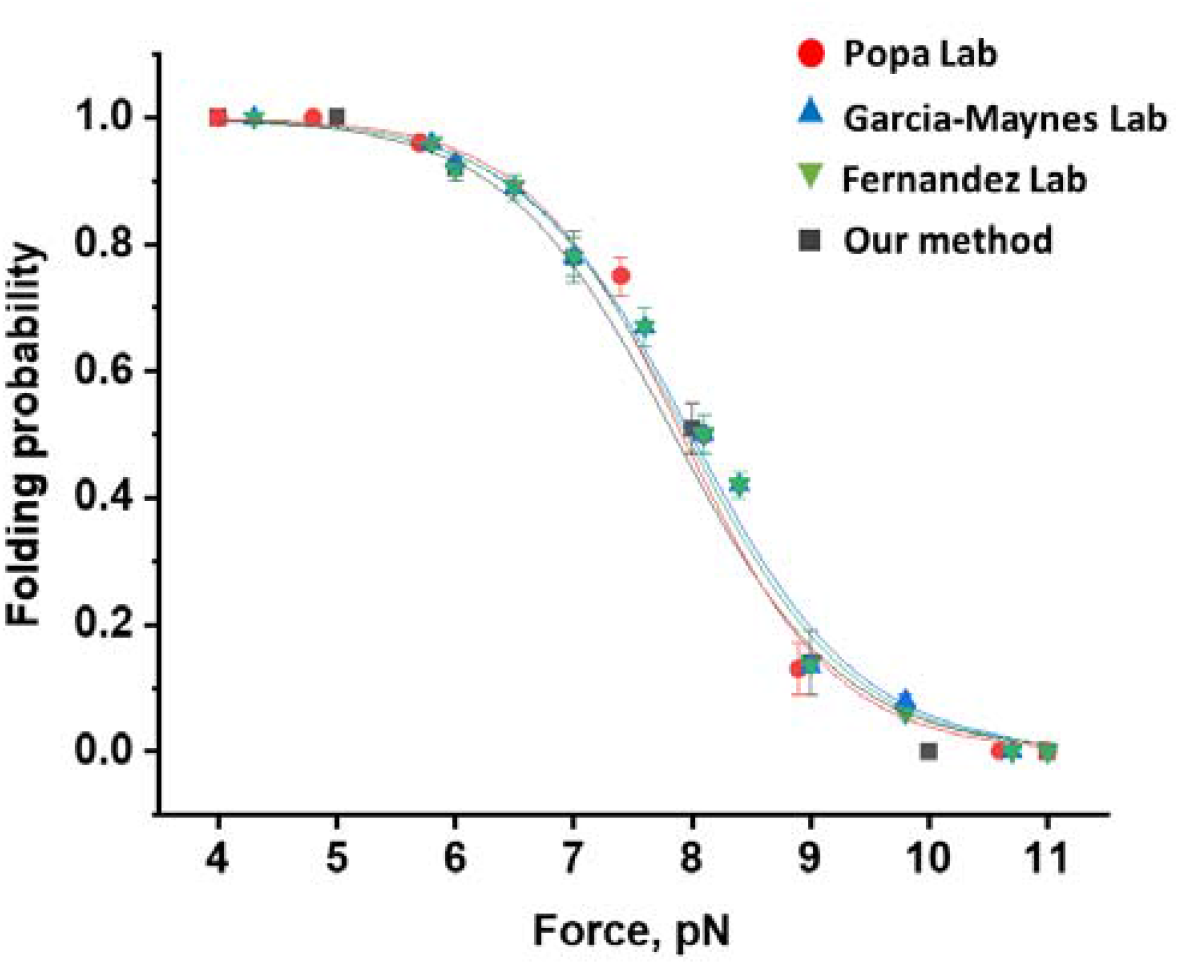
Confirmation of the force calibration by comparing folding probability of protein L: We have analyzed the folding probability by our force calibration method and compared with the folding probability values of protein L, studied by other groups.

### Chaperones interaction with the folded/collapsed state of protein L

To check whether the chaperones interact with the folded/collapsed state of protein L substrate, we linearly increased the force from 4 to 80 pN at a loading rate and monitored the unfolding forces of eight domains in the protein L octamer construct (Supplementary Figure 13A). Average unfolding force of protein L domains has been found to decrease with the unfolding force increases in the presence of foldase chaperone (DsbA). This chaperone-altered unfolding force certainly signifies these chaperones interacts with the folded state of protein L (Supplementary Figure 13B). We also reconciled our observation using talin domain, where similar changes in the average unfolding force has been observed (Supplementary Figure 13C).

**Supplementary Figure 13:**
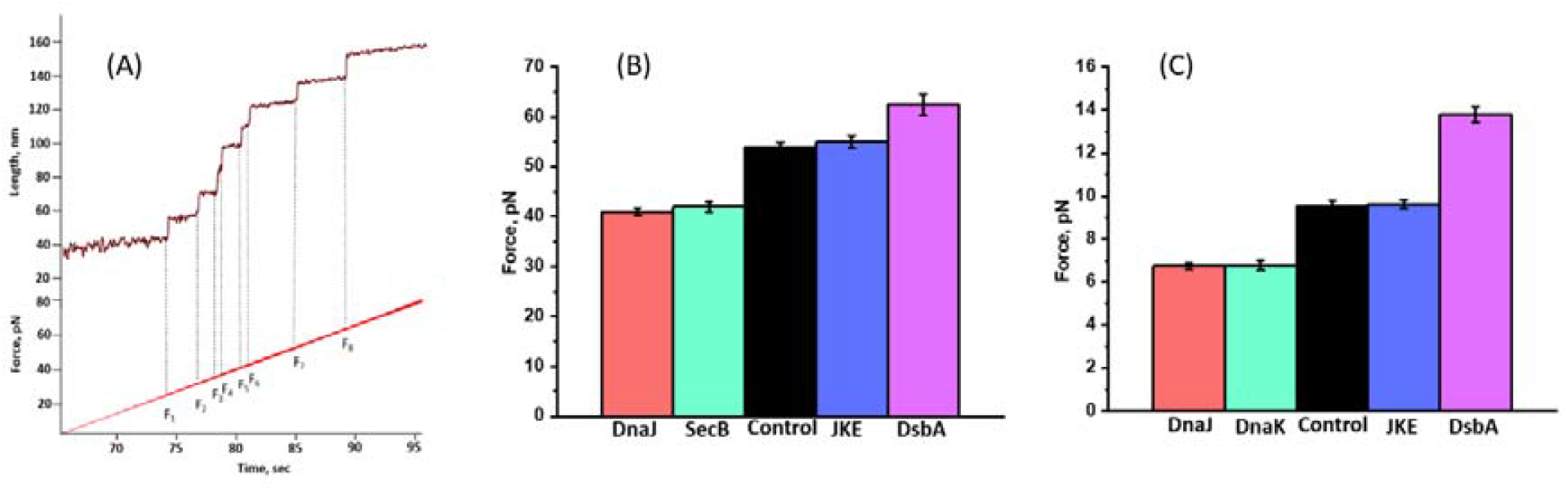
(A) Force ramp study of protein L: We increased the force linearly at a loading rate to monitor the unfolding forces of eight domains in the protein L octamer construct. **(B) Unfolding force of protein L:** the average unfolding force of protein L domains has been found to decrease in the presence of different unfoldase chaperones such as DnaJ, SecB than in their absence (control), while the unfolding force increases in the presence of foldase chaperone (DsbA). This chaperone-altered unfolding force certainly signifies these chaperones interacts with the folded state of protein L. Error bars are s.e.m. **(C) Unfolding force of talin:** Similarly, chaperones also modulate the average unfolding force of talin: decreases with unfoldase chaperones, while increases with the foldase chaperones. However, in the presence of DnaKJE chaperone complex they are not able to change the unfolding force.

**Supplementary Figure 14:**
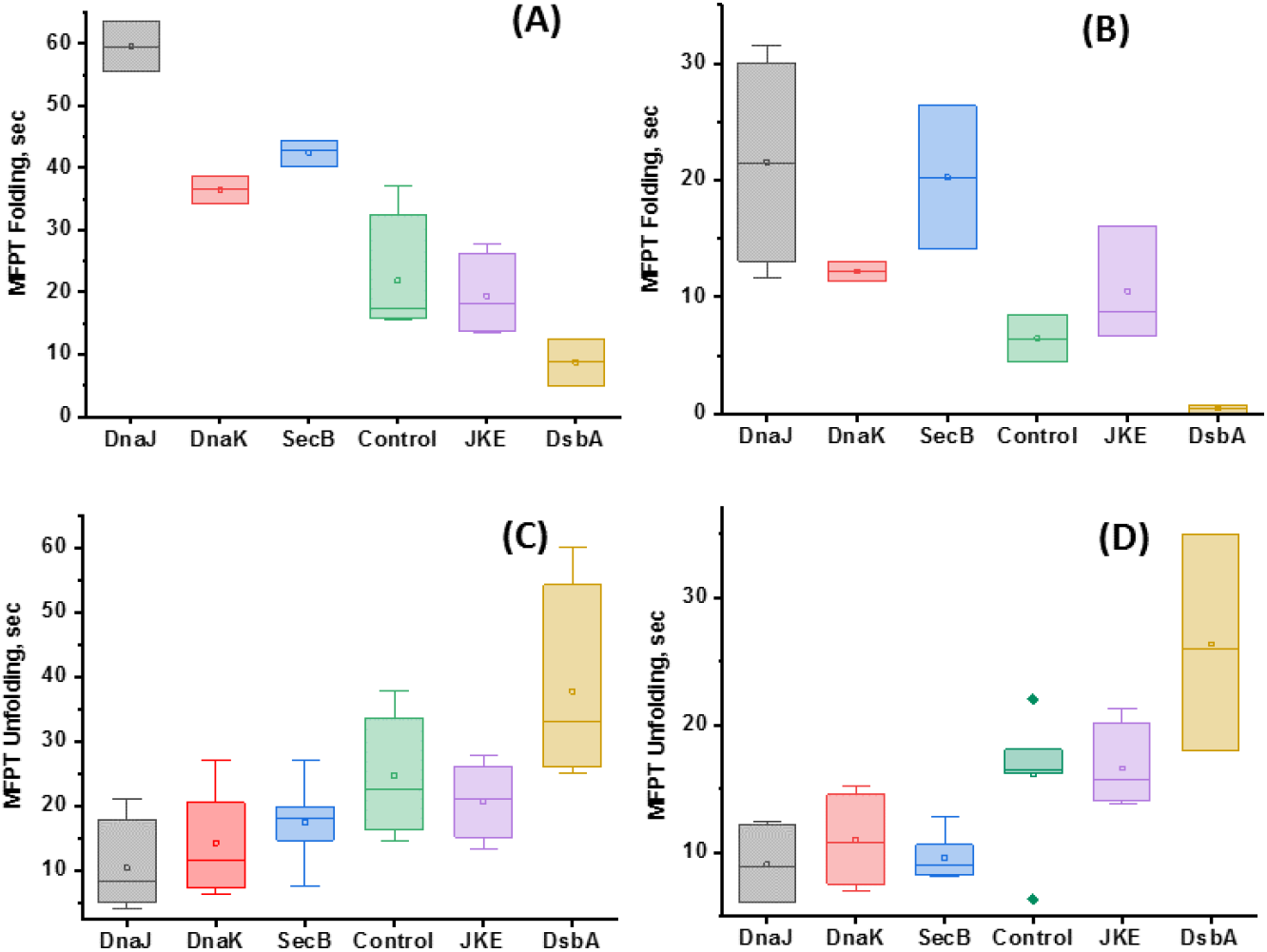
ANOVA (one way) analysis of the refolding and unfolding MFPT values: MFPT refolding values are plotted for different chaperones at two different forces: 7 pN (A) and 6 pN (B). Similarly, MFPT unfolding values are plotted at 45 pN (C) and 50 pN (D) to perform their ANOVA analysis. The analysis shows that in each of the cases, the population means (here refolding and unfolding MFPT values in sec with different chaperones) are statistically different. For example, at 7 pN force, the R-sq.= is 0.88 for MFPT refolding and for MFPT unfolding at 50 pN, R-sq. = 0.86, implying that these MFPT times are statistically significant at p=0.05 level.

**Supplementary Figures 15:**
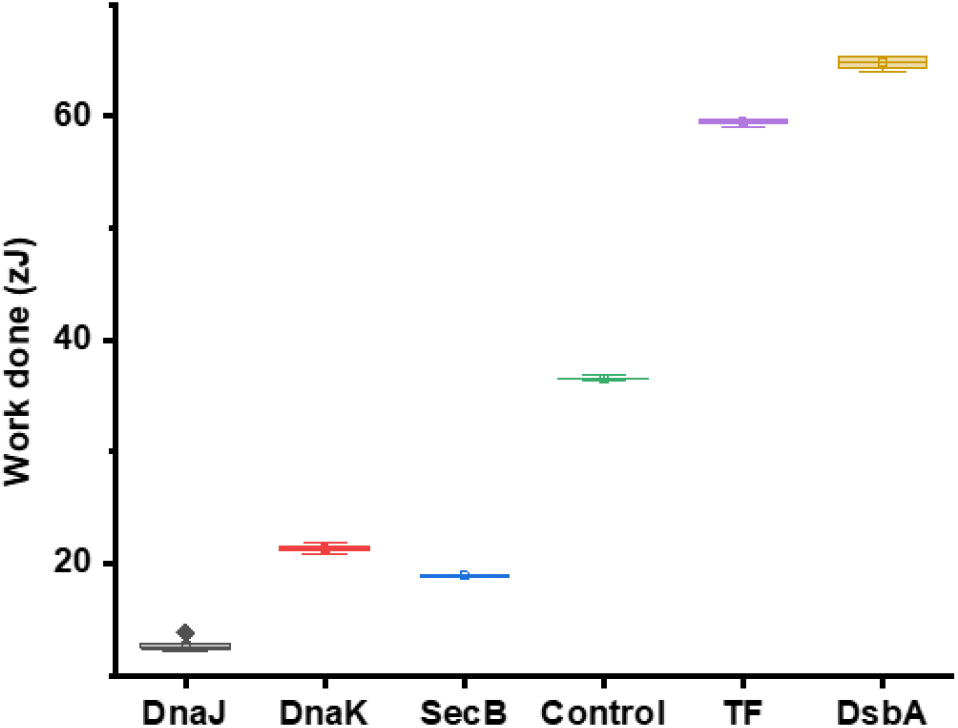
ANOVA analysis of the mechanical work done of protein L folding in the presence of different chaperones: ANOVA analysis shows the population data i.e. work done values are statistically different at p=0.05 level and the R square value is 0.99.

